# The infinitesimal model

**DOI:** 10.1101/039768

**Authors:** N H Barton, A M Etheridge, A Véber

## Abstract

Our focus here is on the *infinitesimal model*. In this model, one or several quantitative traits are described as the sum of a genetic and a non-genetic component, the first being distributed as a normal random variable centred at the average of the parental genetic components, and with a variance independent of the parental traits. We first review the long history of the infinitesimal model in quantitative genetics. Then we provide a definition of the model at the phenotypic level in terms of individual trait values and relationships between individuals, but including different evolutionary processes: genetic drift, recombination, selection, mutation, population structure, … We give a range of examples of its application to evolutionary questions related to stabilising selection, assortative mating, effective population size and response to selection, habitat preference and speciation. We provide a mathematical justification of the model as the limit as the number *M* of underlying loci tends to infinity of a model with Mendelian inheritance, mutation and environmental noise, when the genetic component of the trait is purely additive. We also show how the model generalises to include epistatic effects. In each case, by conditioning on the pedigree relating individuals in the population, we incorporate arbitrary selection and population structure. We suppose that we can observe the pedigree up to the present generation, together with all the ancestral traits, and we show, in particular, that the genetic components of the individual trait values in the current generation are indeed normally distributed with a variance independent of ancestral traits, up to an error of order 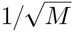. Simulations suggest that in particular cases the convergence may be as fast as 1/*M*.

## 1 Introduction

A simple and robust model for the inheritance of quantitative traits is that these are the sum of a genetic and a non-genetic (environmental) component, and that the genetic component of offspring traits follows a normal distribution around the average of the parents; this distribution has a variance that is independent of the parental values, and, in a large outcrossing population, remains constant despite selection.

This model has its roots in the observations of Galton (1877, 1885, 1889), and their analysis by Pearson (1896, 1897). Fisher (1918) showed that trait values and their (co)variances can be broken down into components, and that the phenotypic observation of constant within-family variance is consistent with a large number of Mendelian factors, with additive effects. This limiting model has become known as the *infinitesimal model*, and can be extended to include all the evolutionary processes: recombination, mutation, random sampling drift, migration and selection. However, although it has long been central to practical breeding, where it forms the genetic basis for the *animal model*, it is relatively little used in evolutionary modelling (see Kruuk 2004 and Hill & Kirkpatrick 2010 for a review). Moreover, there seems to be no agreement on what precisely is meant by the infinitesimal model, nor on the conditions under which it is expected to apply.

This paper provides a concise summary of the model, and shows that it can be accurate even with epistasis. The reason can be understood intuitively, as follows. The classical theory of quantitative genetics gives a remarkably general description of neutral evolution, in which the covariance in the values of a trait across individuals is a linear combination of a set of variance components, with coefficients determined by the probability of identity of sets of genes. Selection rapidly changes the trait mean, at a rate proportional to the additive genetic variance. However, when the trait depends on large numbers of genes, each of which makes a small contribution, selection has a negligible effect on the variance contributed by any individual locus. At the individual level, conditioning on the trait value hardly alters the distribution of effects of any one gene, at least in the short term; therefore, this distribution can be assumed constant. Importantly, it is not that allele frequencies do not change under the infinitesimal model: allele frequencies may change substantially due to random drift, mutation and migration; the key assumption is that selection only slightly perturbs the neutral distribution at any particular locus (Fisher 1918; Robertson 1960).

For neutral populations, the validity of the infinitesimal model is well known. Our results here incorporate selection, mutation, random drift, population structure and epistasis. Since selection acts on traits, and thus distorts the pedigree, by conditioning on the pedigree and the trait values in all previous generations, we incorporate arbitrary forms of selection. In the same way, we capture population structure. We prove that the distribution of traits in the population is a multivariate normal distribution in which, although selection may shift the mean, covariance is determined entirely by the pedigree and is independent of trait values. Conditioning on knowing just some of the trait values in the pedigree shifts the mean trait values in other families by a linear functional of the conditioned values, but the variances within families are unaltered.

After outlining the history of the infinitesimal model, we define it directly as a model for the distribution of phenotypes in a population. Initially, we implicitly assume an additive trait, but include all evolutionary processes. For simplicity, throughout, we shall assume no linkage. We shall explain how to simulate the model, both at the level of the population and the individual, and outline some of its applications. We then derive the infinitesimal model as a limit of a model of Mendelian inheritance, showing the conditions under which it is accurate. Finally, we show how the infinitesimal model extends to allow for epistasis, before presenting simulations that illustrate the main results.

## 2 The classical model

### 2.1 History

Although the infinitesimal model is named for its justification as the limit of infinitely many Mendelian genes, it can be defined purely phenotypically, and its origins trace back well before the rediscovery of Mendel’s work in 1900.

In one of the earliest quantitative discussions of heredity, Fleeming Jenkin (1867) argued that blending inheritance could have no effect in the long term: a white man stranded on a tropical island would leave offspring who, over successive generations, would approach ever closer to the dark-skinned native population. Davis (1871) pointed out that in a large and stable population, an individual is expected to leave two children, four grandchildren, and so on, so that his total expected contribution is constant through time. Nevertheless, if offspring are precisely intermediate between their parents, the range of variation in the population must necessarily decrease. Darwin saw this as a serious problem for his theory, which required a source of variation to counter blending inheritance. (See Bulmer, 2004, for a detailed discussion of Jenkin’s argument).

Francis Galton gathered extensive data on the inheritance of continuous traits, and introduced many ideas that are now central to quantitative genetics. In experiments with sweet peas, he showed that seeds of offspring grown from seeds of different weights followed a normal distribution with a mean that reverted towards the population mean, and with variance independent of the parents’ weight: “I was certainly astonished to find the family variability of the produce of the little seeds to be equal to that of the big ones, but so it was, and I thankfully accept the fact, for if it had been otherwise, I cannot imagine, from theoretical considerations, how the problem could be solved” (Galton, 1877, p.513). (In Galton’s experiments with sweet peas, plants were self-fertilised, so that the variance in the population is, in fact, expected to decrease.) He saw a similar pattern for human height, and showed that the joint distribution of offspring and mid-parent is bivariate normal (Galton, 1885). Moreover, he understood that the variance of the population could remain stable under the joint influence of random mating, reversion of offspring towards the population mean, and generation of variance amongst offspring. Galton (1877) calculated the equilibrium variance, allowing for Gaussian stabilising selection, a calculation next made by Bulmer (1971) and Cavalli-Sforza & Bodmer (1971), nearly a century later.

Galton (1885, 1889) tried to explain his observations by formulating his ‘law of ancestral heredity’, which divided an individual’s phenotype into geometrically declining contributions from parents, grandparents, great-grandparents, …; he interpreted this contribution from distant ancestors as being due to inherited factors which have some probability, *p*, of being expressed in each generation. Bulmer (1998) shows that Galton’s law is equivalent to the quantitative genetics of an additive trait, with *p* being replaced by the heritability, *h*^2^ = *V*_*A*_/*V*_*P*_ (where *V*_*P*_ is the total phe-notypic variance and *V*_*A*_ the additive genetic variance of the trait); however, *h*^2^ may vary from trait to trait, whereas Galton assumed that it is a constant parameter of the mechanism of inheritance. Galton’s model explains reversion of offspring towards the population mean as being due to expression of factors inherited from earlier generations. In contrast, under Mendelian inheritance, reversion to the mean arises because selection acts on the phenotypic variance, *V*_*P*_, whereas only additive genetic variation, *V*_*A*_, is passed on; the deviation of offspring is therefore *h*^2^ = *V*_*A*_/*V*^*P*^ times that of the selected parents. Pearson (1896, 1897) introduced matrix algebra and multiple regression, to put Galton’s ancestral law on a firm mathematical basis. However, he treated the problem as the statistical description of a population of relatives, rather than following Galton in devising a mechanistic explanation of heredity (Magnello, 1998).

After 1900, there was a bitter dispute between those studying the Mendelian inheritance of discrete characters, and those biometricians who studied the inheritance of continuous traits (Provine, 1971). Pearson (1904a, 1904b, 1909) understood that Mendelian factors could account for continuous variation, but found that if there were complete dominance, correlations between relatives did not agree with observations (see Magnello, 1998). Yule (1902, 1906) showed that if incomplete dominance and random ‘environmental’ variation are included, then arbitrary correlations could be explained; however, these ideas were not developed further until Fisher’s definitive (1918) paper.

During the following years, quantitative genetics developed quite separately from the population genetics of discrete genes. Fisher and Wright established the basic theory for correlation between relatives and for the effects of inbreeding, Wright was involved in practical animal breeding, and Haldane (1931) showed how selection on a trait affects the constituent alleles. However, the bulk of their work was on the evolution of single loci, and even the basic theory for the response of continuous traits to selection developed slowly. The change over one generation is implicit in Galton’s regression of offspring on mid-parent, and the multivariate version is given by Pearson (1896). However, the classic ‘breeders’ equation’ was only written in its modern form by Lush (1937); see Hill & Kirkpatrick (2010). Fisher’s (1918) analysis of genetic variance was developed into a sophisticated theory in the 1950’s (e.g. Henderson, 1953; Cockerham, 1954; Kempthorne, 1954; see Hill, 2014), but this did not become widely known. Quantitative genetics came back into contact with evolutionary questions through Robertson (1966), who formulated the ‘secondary theorem’, which states that the rate of change of a trait’s mean due to selection equals its covariance with relative fitness. Robertson (1960) also showed that under the infinitesimal model, the ultimate response to selection equals the response in the first generation, multiplied by twice the effective population size; he showed that this can be understood through the increase in fixation probability of individual alleles caused by weak selection (*N s* ≪ 1). (We discuss this in more detail when we discuss applications of the infinitesimal model in Section 2.3 below.) Bulmer (1971) investigated the effect of stabilising selection and mutation on trait variance. Lande (1976a,b) investigated the long-term evolution of multivariate traits, extending Wright’s ‘adaptive landscape’ to quantitative traits, and Lande & Arnold (1983) showed how multivariate selection can be measured in natural populations. It is striking that though these methods trace back to Galton and Pearson, they did not become widely used in evolutionary biology for more than 70 years. Indeed, the sophisticated ‘animal model’, widely used in animal breeding, has only been applied to analyse natural populations over the past 15 years (Kruuk, 2004).

Despite the revival of interest (both theoretical and empirical) in ‘evolutionary quantitative genetics’ in recent decades, the infinitesimal model itself has received little attention. Bulmer (1971) showed that assuming a large number of unlinked loci with additive effects, the joint distribution of a set of relatives is multivariate normal, conditional on the parents; Lange (1978) gave a more detailed derivation. Again assuming additivity, Dawson (1997) showed that certain kinds of linkage disequilibrium could cause the distribution amongst offspring to depend on the parental values. Bulmer (1974) and Santiago (1998) extended the infinitesimal model to allow for linkage. Turelli & Barton (1994) gave a general treatment of epistasis, which allows for linkage and multiple alleles. They showed that provided that *k*th order epistatic coefficients scale correctly with the number of loci, *M*, then the effect of selection on the trait depends only on the variance of effects at each locus, and the linkage disequilibria between them. The additive genetic variance will change slowly under selection, and can be assumed constant for 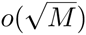 generations. However, their treatment did not include mutation, population structure, or random drift.

### 2.2 Definition of the phenotypic model

We begin by defining the infinitesimal model in terms of the phenotypic distribution. In Section 3.1, we derive it as the limit of a large number of Mendelian alleles with additive effects, and that underlying additivity will be implicit in our discussion in this section. However, in Section 3.2 we show that under some conditions the model can be extended to include epistasis and the phenotypic model will, just as in the classical case which we now describe, be determined by systems of recursions for the segregation variance between siblings.

For simplicity, in this section, we ignore non-genetic contributions to the trait; an environmental noise will be explicitly incorporated in our derivations in Section 3.1. We also consider a single trait, but it is straightforward to extend the analysis to multiple traits.

#### The basic model

Consider first the simplest case, of a purely additive trait in a large outcrossing population. Then, the infinitesimal model states that offspring follow a Gaussian distribution around the mean of the two parents, with a variance *V*_0_, that is constant, independent of the parents’ values. With random mating, the population as a whole rapidly converges to a Gaussian with variance 2*V*_0_. To see this, note that if the variance in the parental population is *V*_1_, then that of the mean of two parents sampled at random is *V*1/2, and so that of the offspring generation is *V*1/2 + *V*_0_: at equilibrium, *V_1_ = 2V_0_;* that is half the variance is between families, and half within them. Selection can readily generate arbitrary non-Gaussian distributions: we are free to choose any distribution of phenotypes. However, in the absence of selection such distributions rapidly relax back to a Gaussian with variance 2*V*_0_; the *k*th order cumulants decay by a factor 2^1-*k*^ per generation, for *k* ≥ 3 (Bulmer, 1980).

#### Haploids versus diploids

In this simplest case, it makes no difference whether we follow haploids or diploids. However, the distinction becomes evident when we consider inbreeding and random drift. We can choose to follow haploid individuals, which mate to produce diploids that immediately undergo meiosis to produce the next haploid generation. Alternatively, we can follow diploid individuals, which produce haploid gametes via meiosis, and then immediately fuse to produce the next diploid generation. This results in two distinct approaches to modelling, both of which we describe below.

With no selection, whether we track haploids or diploids makes no fundamental difference. However, when we select, we condition the individual’s full genotype on the value of a polygenic trait; it is then clearly important whether we measure the trait at the haploid or the diploid stage. In principle, selection could act at both stages, but we do not consider this complication. For simplicity, in our derivation in Section 3 we concentrate on the haploid case.

#### Identity by descent and the segregation variance

Variation between siblings is generated by the random segregation of genes from the parental genotypes. To the extent that the genomes that come together in meiosis are related, this segregation variance will be reduced in proportion to the fraction of genes that they share. Imagine an ancestral population, whose members are all unrelated. We suppose that after one round of reproduction all families segregate with variance *V*_0_. The current population descends from this reference population via an arbitrary pedigree. The relation between haploid individuals *i, j* is described by *F*_*i,j*_, the probability that homologous genes descend from the same ancestral gene - that is, are identical by descent from the reference population. Since we are ignoring linkage, and the trait is additive, the variance amongst the haploid offspring from haploid parents *i, j*, is just *V*_0_(1 - *F*_*i,j*_).

For diploids, *F*_*i,j*_ is defined to be the probability of identity between two genes, one from *i*, and one from *j;* when *i = j, F*_*i,i*_ is defined to be the probability of identity by descent of two *distinct* genes in the diploid individual *i*. Meiosis in *i* generates segregation variance proportional to p1 - *F*_*i,i*_). The value of an additive trait in a diploid is the sum of equal contributions from each haploid gamete, and so the segregation variance is *V*_0_ (1 - (*F*_*i,i*_ + *F*_*j,j*_)/2). To see this, one can note that segregation occurs independently to create the 2 parental gametes and, for each of them, conditional on not being identical by descent the ancestral genes are two independent samples from the initial population with variance *V*_0_. This yields an expression for the segregation variance of

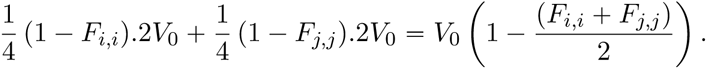

We have defined the infinitesimal model in terms of a constant genetic variance, *V*_0_, in a reference population, together with a matrix of identity by descent. The entries in the matrix increase over time, until they are all one, and all variation is lost. However, we could instead follow the matrix of segregation variance between pairs of individuals. This process evolves with time, but it is Markovian (i.e., depends only on the current state), and it has the advantage that it does not require that we define an ancestral reference population at some arbitrary time in the past. As we shall see below, when we derive the infinitesimal model as a limit of a model of Mendelian inheritance, it is also convenient when we introduce mutation. For haploids, we define *C*_*i,j*_ as the variance amongst offspring from two haploid parents, *C*_*i,j*_ = *V*_0_ (1 - *F*_*i,j*_). For diploids, as we saw above, the variance between siblings depends only on the identity between distinct genes in the parents, and not on the relationship between the two diploid parents. We define *C*_*i,j*_ = *V*_0_(1- *F*_*i,j*_), just as for haploids, but now, the variance amongst the diploid offspring of diploid individuals *i, j* is (*C*_*i,i*_ + *C*_*j,j*_)/2.

Although *F*_*i,j*_ is defined through the probability of identity by descent of discrete genes, it can (in principle) be measured purely through phenotypic measurements of the variance amongst offspring; this is perhaps clearer if we work with the *C*_*i,j*_. The classical infinitesimal model is based on the assumption that the *C*_*i,j*_ depend only on the pedigree relationship between *i* and *j*, and are independent of which trait is measured (to within the constant factor *V*_0_), and of the trait values of the parents. In general, we think of a trait as being the sum of a genotypic value and an environmental deviation, independent of the underlying genetic values. We shall explicitly incorporate this non-genetic variation when we derive the infinitesimal model as a limit of Mendelian inheritance in Section 3. For the moment, we assume additivity and ignore environmental variation, so that the trait value is equal to the genotypic value, which in turn equals the *breeding value*. The breeding value of an individual is defined to be twice the mean deviation from the average phenotypic value, when it is crossed with a randomly chosen individual. The genotypic value can, in principle, be measured as the mean of large numbers of cloned individuals, and the breeding value can be measured through the mean of offspring from crosses made with randomly chosen mates. So, the infinitesimal model can be defined without identifying any specific genes.

#### Recursions for identity by descent

In a randomly mating population of *N* haploid individuals, reproducing under the Wright-Fisher model, the expected identity is *F* =1 - (1 - 1*/N*)^*t*^ after *t* generations. However, we consider the general case, where *F*_*i,j*_ may vary arbitrarily between pairs. For haploids, *F*_*i,i*_ = 1 by definition. The recursion for *F* can be written in terms of a *pedigree matrix, P*_*i,k*_(*t*), which gives the probability that a gene in *i* in generation *t* came from parent *k* in the generation (*t* - 1); each row has two non-zero entries each with value 1/2, unless the individual is produced by selfing, in which case there is a single entry with value 1. Thus,

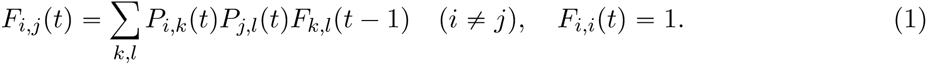

For diploids, the corresponding recursion for *F* is

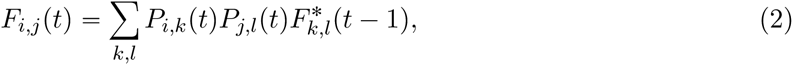

where

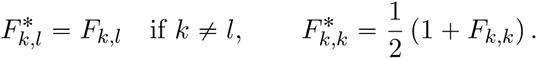

The quantity 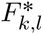 is the probability of identity of two genes drawn *independently* from *k, l;* if *k = l*, then the probability of drawing the same gene twice is one half.

If we work with the segregation variances, *C*_*i,j*_, then the recursion for haploids is

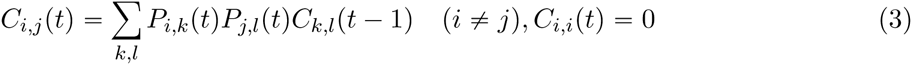

and for diploids is

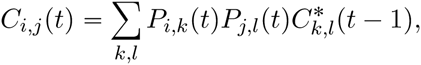

where

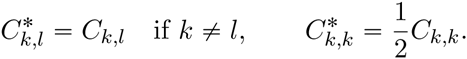

Note that the variance in the base population, *V*_0_, does not appear explicitly: the future dynamics are entirely determined by the variation that is released through recombination between any pair of genomes. Although the precise recursions that we have written down are particular to the additive model, analogous recursions characterise the segregation variance in our more complex models that incorporate house of cards mutation and epistasis. The key fact will be that it is the pedigree relatedness between individuals that drives the recursions. As long as the variances in the parental population are sufficiently large relative to the effect of individual alleles, knowing the trait values of the parents has a negligible effect on the segregation variance; in other words, the infinitesimal model remains valid.

#### Simulating the infinitesimal

The infinitesimal model can be simulated either at the level of the individual, or the population. An individual-based simulation must follow the breeding values of each individual, *z*_*i*_, and the relatedness between individuals, *F*_*i,j*_. Extension to multiple traits would require that we follow vectors of breeding values. Since the main computational effort is in calculating the matrix of identities, this is not much extra burden. The matrix of identities can be iterated efficiently by representing Equations (1) and (2) in matrix form, but the size of the population is ultimately limited by the memory needed to store *F*_*i,j*_. However, in large populations *F*_*i,j*_ typically approaches the same small value between almost all pairs; thus, it can be approximated as a constant plus a sparse matrix that tracks close relatives. Populations of many thousands can then be simulated (e.g. Barton and Etheridge, 2011).

Provided that the pedigree determined by the matrix *F*_*i,j*_ is not too skewed towards large contributions from particular individuals, then we can also simulate very large populations by following the distribution of the trait and the *average* value of *F*_*i,j*_ through time. To do this, first, the trait distribution must be approximated by a discrete vector; selection on the trait is represented by multiplying the trait distribution by the fitness. Since reproduction involves a convolution between the parents’ distributions and the Gaussian distribution of offspring, it is convenient to follow the (discrete) Fourier transform: convolution of distributions corresponds to multiplication of their transforms. In each generation, there must be a conversion between the distribution and its transform, which can be done efficiently using the fast Fourier transform algorithm (Gauss, 1866; Cooley and Tukey, 1965). Evidently, the approximation that all individuals are related by the same *F*_*i,j*_ will not always be realistic, in which case an individual based approach becomes essential.

#### Mutation

Pragmatically, for traits determined by a very large number of loci, mutation can be included by scaling the recursion to account for alleles that are replaced by mutants and adding a constant, which may depend on *t*, to every element of the matrix *C*^*i,j*^ in each generation to account for the variance introduced by mutation (Wray, 1990), see equation (5) below. Mutation may be biased: in particular, we expect mutation to decrease traits that have been under directional selection, and so to decrease fitness. This can be described by scaling the mean of the offspring, by a constant (1 - *μ*) say, and shifting it by another constant *δ*_*μ*_. Under this extension to the infinitesimal model, *μ* is assumed constant and the distribution of offspring, conditional on their parents, is assumed Gaussian.

In Section 3.1, we obtain this as the limit of a Mendelian model of an additive trait with mutation at every locus governed by the *House of Cards* approximation (Turelli, 1984). Mutation is assumed to generate new alleles with random effects that are independent of the original values: with probability (1-*μ*_*l*_), the allelic effect at locus *l* is inherited from the parent, whilst with probability *μ*_*l*_ it is drawn from a constant distribution with mean *θ*_*μ,l*_ and variance 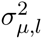. In fact, for mathematical convenience, we shall take *μ*_*l*_ to be constant across loci, but the allelic effects will be allowed to be locus-dependent. The recursion for haploids then becomes

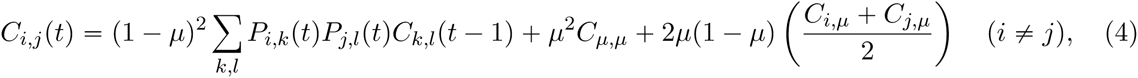

where formally *C*_*i,μ*_ is the segregation variance in a family in which one parent is sampled from the family with label *i* (with no mutations) and the other is mutant at every locus, and *C*_*μ,μ*_ is the segregation variance in a family in which both parents carry mutant alleles at all loci. Of course such purely mutant individuals never appear in the population as, for large *M*, only a fraction *μ* of an individual’s loci are affected by a mutation. One should instead see the parameters *C*_*i,μ*_ and *C*_*μ,μ*_ as summing the contributions of every locus to the segregation variance under the different possible scenarii. For every given locus, the case when one parental allelic state is transmitted without mutation and the other parental allelic state is a random draw from the mutant allele distribution (which occurs with probability 2*μ*(1 - *μ*)) adds a contribution to the variance which appears in *C*_*i,μ*_ or *C*_*j,μ*_ (depending on which parent transmits its allele without mutation); on the other hand, the contribution to the variance of the case when both parental allelic states are mutants (which happens with probability *μ*^2^) appears by definition in *C*_*μ,μ*_.

Since a given allelic effect in individual *i* is drawn from the distribution of allelic effects in the base population with probability (1 -*μ*)^*t-*1^, or is sampled from the mutant distribution otherwise, equation (4) can be reexpressed as

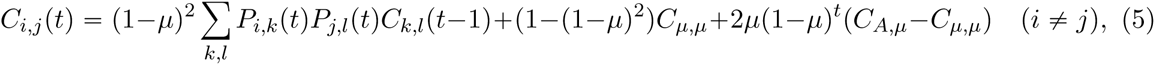

where *C*_*A,μ*_ is the segregation variance in a family in which one parent is purely ancestral (no mutations) and the other is mutant at every locus. It is in this form that it is proved in Section 3.1. Note that if *i = j*, then the first term is zero.

When the per-locus mutation probability *μ* is very small, as will usually be the case, equation (5) simplifies into

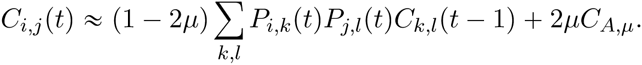

An expression for *C*_*A,μ*_ in terms of ancestral and mutated allelic effects can be found in equation (11). Assuming that 2*μC*_*A,μ*_ ≈ *V*_*m*_/2 > 0 when *μ* is small, we arrive at

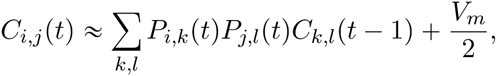

where *V*_*m*_ can be interpreted as the total increase in additive genetic variance in the population due to mutation per generation.

#### Population structure and gene flow

When defined in terms of individual trait values and relationships, the infinitesimal model automatically incorporates arbitrary population structure. A well-mixed reference population may split into separate demes, so that the probability of identity *F* between genes from within the same population would increase relative to that between populations. If the demic structure were permanent, then it would be more natural simply to follow the segregation variance *C*^*i,j*^, which would be higher from crosses between populations than within them. If, for whatever reason, the sub-populations in different demes diverge, then the distribution of trait values within a single deme will not be Gaussian, even though, under the infinitesimal model, the distribution amongst offspring of given parents will always be Gaussian. The same is true for any pedigree which is not ‘well-mixed’; the key point is that parental trait values determine the mean trait value among offspring, but the segregation variances that determine the variance of offspring traits are completely determined by the pedigree.

The power of the infinitesimal model in capturing population structure comes at a price; for a large population it may not be practicable to trace the breeding values and relatedness of all individuals. In that case, one could try to approximate the infinitesimal model, for example by assuming that the trait distribution within populations is approximately Gaussian, or that relationships (or equivalently, segregation variance) between individuals within subpopulations are homogeneous. Such approximations may be delicate, since they might need to take account of the reduction in effective migration rate, and the increase in the rate of random drift, due to selection. However, it is important to realise that the infinitesimal model itself can be defined at the level of individuals, even when trait distributions are far from Gaussian, and relationships are heterogeneous.

#### Multivariate normality

Even though the trait distribution across the whole population may be far from Gaussian, the multivariate normal will play a central role in our analysis. The deviations of individual trait values from their mid-parent value will be described by a multivariate normal whose variance-covariance matrix is independent of the parental traits. We establish this result conditional on knowing the pedigree and all ancestral traits, but equally we could have conditioned on the values of any subset of relatives and the same would hold true: expected trait values would be a linear functional (determined by the pedigree) of the values on which we conditioned, but segregation variances would be unchanged.

Bulmer (1971) and Lange (1978) showed that the unconditional joint distribution of traits converges to a multivariate normal. Lange also allows for some linkage among loci by allowing inheritance to be dependent among loci at distance at most *q* from one another. We do not include linkage in our analysis, although as long as recombination is sufficiently fast, our results should hold true, but whereas the rate of convergence to the infinitesimal model when we have unlinked loci is 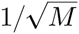, with linkage it will be 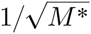, where *M*\* is an ‘effective’ number of loci. Bulmer assumed random mating, while Lange’s proof is for individuals related through a given pedigree. However, as Lange remarks, his result gives no control on the rate of convergence. This is essential if we wish to approximate the conditional distribution, knowing some ancestral trait values. It is also needed in assessing the accuracy of the infinitesimal model as an approximation.

#### Epistasis

Thus far, we have defined the infinitesimal model for the additive case. Evidently we cannot extend it to arbitrary epistatic interactions. For example, if *Z* is a purely additive trait, then *Z*^2^ is a sum of additive and pairwise epistatic components. Since the square of a normally distributed random variable is not normally distributed, the infinitesimal model must clearly break down. However, under some conditions (which we lay out in Section 3.2), even though there can be significant variance due to epistatic components, the infinitesimal model still holds with epistasis.

In general, with epistasis the individual phenotype is

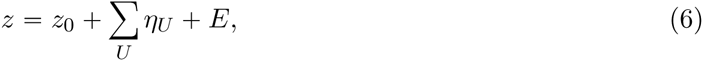

where the sum is over the average effects *η*_*U*_ of all sets *U* of distinct loci, and *E* is a random non-genetic component that is assumed to have a distribution independent of genotype, and independent between individuals (see e.g. Chapter 7 in Lynch & Walsh, 1997). The sets of genes *U* descend from a homologous set in the base population, which in general will be scattered over many individuals. The *η*_*U*_ are defined as the marginal effects of the set of genes *U*, that remain after accounting for the effects of all subsets of *U*. If the base population is in linkage equilibrium, then the *η*_*U*_ are uncorrelated. The sum of the variances contributed by sets of size |*U*| = *k* is the *k*th order epistatic variance, Σ_|*U*|=*k*_*V*_*U*_ = *V*_*A*(*k*)_. In contrast to the additive case, we see correlations between the deviations ∆*Z* from the mid-parental trait values of distinct individuals. The covariance between two distinct individuals is

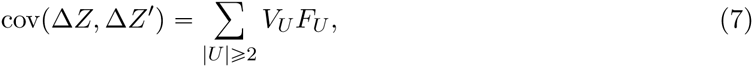

where *F*_*U*_ is the probability that the set of genes *U* in the two individuals are all identical by descent. If loci are unlinked, then this depends only on the number of genes in the set, so that *F*_*U*_ = *F*_*k*_ where *k* = |*U*|; *F*_*k*_ is given by a recursion on the pedigree similar to that described above for pairwise identities (corresponding to *F*_1_). It is more complicated, because we need to track the probability of identity for genes in up to 2*k* individuals. However, if identity at different loci is uncorrelated, then 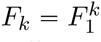, where *F*_1_ is the pairwise recursion defined above. Unless inbreeding is intense, this is typically a good approximation (Barton and Turelli, 2004).

This analysis of genetic variance into components applies regardless of the number of loci. Crucially, however, if the joint distribution of the components of trait values across the pedigree is multivariate normal, then the mean and covariance completely define that distribution. In the following, we outline the proof that this is indeed the case, provided that the number of loci with which any particular gene can interact is not too large. We achieve this “sparseness” by partitioning loci into nonoverlapping blocks which are not allowed to interact. However, this condition is motivated by the generalised central limit theorem that we use in the proof and we illustrate with a numerical example in Section 3.3 that the result will hold true with other types of weak epistatic interactions. Our derivation also requires that for each epistatic component, if we condition on the allelic states at all but one of the loci then the average over the distribution at that locus in the ancestral population is zero.

We emphasise that the components of phenotype, *η*_*U*_, and the corresponding variances, *V*^*U*^, are defined relative to the base population. In any particular descendant population, the trait mean will differ as a result of mutation (which we exclude from our analysis), selection and random drift. With epistasis, the effects relative to the new population will be different, and so the variance components defined for this descendant population will also differ. This can lead to the “conversion” of epistatic variance into additive variance (Barton and Turelli, 2004). We do not consider this issue here, since we define variance components relative to the base population. However, it is straightforward to change the reference point.

### 2.3 Applications of the infinitesimal model

We have defined the infinitesimal model in terms of individual trait values and relationships between individuals, without referring explicitly to discrete genes. This is essentially the ‘animal model’, which is the basis for practical animal breeding, though extended to include mutation. In practical applications, the ‘animal model’ is typically applied to a given pedigree, and is used to estimate breeding values and genetic variances conditional on that pedigree (Hill, 2014). In recent years, it has also been applied to parameter estimation in natural populations (Kruuk, 2004). However, it has been surprisingly little used for addressing evolutionary questions. Here, we illustrate the power of the infinitesimal model as a tool for understanding aspects of evolution, by presenting a range of examples related to stabilising selection, assortative mating, effective population size and response to selection, habitat preference and speciation.

Perhaps the simplest non-trivial application of the infinitesimal model is to understand stabilising selection (Galton, 1877; Bulmer, 1971; Lande, 1976a). Suppose that the distribution of a trait in a parental population is Gaussian with mean 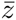 and variance *V*_*g*_. If fitness is a Gaussian function of trait value, with mean *z*_0_ and variance *V*^*s*^, then after selection the new trait distribution is obtained by multiplying the density of a trait by the fitness of that trait and renormalising to have total mass one, resulting in a Gaussian with mean

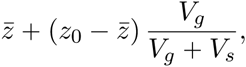

and variance

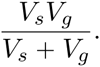

After random mating and reproduction under the infinitesimal model (without mutation), the mean remains the same, and the variance is

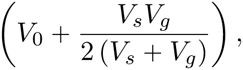

where *V*_0_ is the segregation variance, and 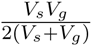 is the variance of the mean of two randomly mated parents. Therefore, at equilibrium, the genetic variance immediately after reproduction is

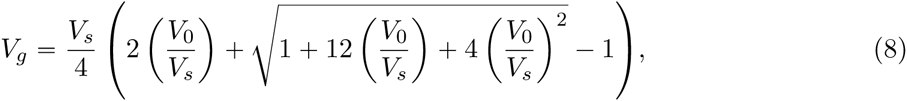

which decreases from 2*V*_*0*_ (the value we obtained for a neutral population) when stabilising selection is weak (*V*_*s*_ ≫ *V*_0_), to *V*_0_ when stabilising selection is so strong as to eliminate all variation (*V*_*s*_ ≪ *V*_0_). The effect of assortative mating is more surprising. Suppose that the relative contribution of pairs with values *z*_1_, *z*_2_ is proportional to

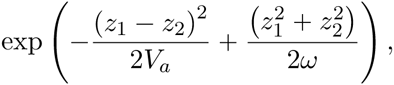

where *ω* is chosen so that there is no direct selection on individuals (see, for example, Appendix 4 of Polechova & Barton 2005 for an expression for the *ω* that achieves this). The mean does not change, but assortment results in a higher variance in the mid-parent value than under random mating. To understand this, note that because the contribution of pairs of individuals is greater for individuals with similar trait values, more extreme traits are less likely to be pulled towards the mean; indeed in the most extreme case, individuals would only reproduce with others with an identical trait value and so the distribution of mid-parent values would have the same variance as that of the whole parental population. Provided that assortment is not too strong *(V*_*a*_ > 4*V*_0_), there is an equilibrium genetic variance 2*V*_0_ (*V*_*a*_ - 2*V*_0_) / (*V*_*a*_ - 4*V*_0_). However, if assortment is very strong, *(V*_*a*_ < 4*V*_0_), the variance increases without limit. In reality, either the infinitesimal model would break down as genetic limits to the trait are approached, or stabilising selection would prevent indefinite divergence.

This simple model makes the important point that assortative mating alone can lead to indefinite divergence, and, ultimately, speciation (Polechova & Barton, 2005). The infinitesimal model can also be extended to model the joint evolution of habitat preference and viability in two different niches, both being represented as continuous traits (Barton, 2010). Assortative mating, and eventual reproductive isolation, then arise as a by-product of preferences for different habitats, provided mating occurs within those habitats (Diehl & Bush, 1989).

In a random-mating population of effective size *N*_*e*_ of diploid individuals, the segregation variance decreases by a factor (1 - 1/2*N*_*e*_) per generation. The infinitesimal model predicts that the response to steady directional selection will decrease at the same rate and so, summing over generations (a geometric sum), Robertson (1960) found the total response to selection to be just 2*N*^*e*^ times the change in the first generation. He also found an alternative derivation of the same result, by considering the increase in fixation probability of neutral alleles. This alternative argument fails if *N*_*e*_*s* is not small and so, in the process he showed that the validity of the infinitesimal model rests on the assumption that the alleles responsible for trait variation are nearly neutral (i.e., *N*_*e*_*s* small). Hill (1982) extended Robertson’s work to include mutation that introduces genetic variance at a rate *V*_*m*_. The genetic variance then reaches an equilibrium between mutation and random drift, *V*_*g*_ = 2*N*_*e*_*V*_*m*_, and the response to directional selection is proportional to this variance. In large populations, selection will tend to reduce genetic variance that is due to alleles with large *N*_*e*_*s*; however, some component of the genetic variance will be due to more weakly selected alleles with *N*_*e*_*s* small. In a survey of selection experiments, Weber & Diggins (1990) showed that the ratio of the response to selection after fifty generations to that after just one generation is somewhat less than predicted by the infinitesimal model. This suggests that for some alleles, *N*_*e*_*s* is large; nonetheless the response to selection is largely explained by the infinitesimal model.

Selection on heritable traits can greatly inflate the rate of random drift: genes that find themselves in a fit genetic background in one generation will tend to be in a fitter background in subsequent generations, even if all loci are unlinked; this correlation in fitness across generations increases the rate of sampling drift (Robertson, 1961). The infinitesimal model can be used to estimate this inflation, by finding the variance in reproductive value (Barton & Etheridge, 2011), and the decrease in fixation probability of favourable alleles (Weissman & Barton, 2012).

Apart from these few examples, the infinitesimal model has hardly been used in evolutionary modelling. It should not be confused with two other models that have been used more extensively. Kimura (1965) investigated the distribution of effects of alleles at a *single* locus, and approximated this *continuum-of-alleles* model by a Gaussian; Lande (1976a) developed this model to investigate maintenance of variation by mutation, despite stabilizing selection. This is a quite different approach from the infinitesimal model, which requires no strong assumptions about the distribution of effects at each locus, and which does not assume a Gaussian distribution of trait values. A second model that bears a superficial resemblance to the infinitesimal model is the *hypergeometric* or *symmetric* approximation, which assumes that the trait is determined by additive loci of equal effect, and that all genotypes that give the same trait value are equally frequent (Kondrashov, 1984; Doebeli, 1996; Barton & Shpak, 2000). This is a very strong assumption; the symmetry between genotypes may hold under disruptive selection, but is unstable under stabilising selection, when any one of the many optimal genotypes tends to fix (Wright, 1935).

## 3 The infinitesimal model as the limit of Mendelian inheritance

In this section, we turn to a justification of the infinitesimal model as a limit of a model of Mendelian inheritance, when trait values are determined by a large number of Mendelian factors, each of small effect. In the classical setting, traits are additive, but essentially the same arguments that allow us to justify the model for additive traits will also allow us to incorporate ‘house of cards mutation’, and, under suitable conditions, epistasis and (although we don’t spell it out here) dominance.

In all cases, the key issue is this: knowing the segregation variance *V*_0_ in our base population and the pedigree *F* (or, equivalently, the matrices *C* of segregation variances in previous generations), how close is the segregation variance of the offspring of parents *i* and *j* to being independent of the trait values of those parents? It is important to note that this does *not* say that the *pedigree* is independent of the trait value; indeed, for a population undergoing artificial selection, for example, one can expect a strong dependence between trait values and pedigree.

One necessarily expects some dependence of segregation variance on trait values: if the possible trait values are bounded, with a single genotype giving the largest value, say, then meiosis between two copies of this most extreme haploid type, or the products of meiosis from a diploid with the most extreme value, would have zero variance. For any trait values that are close to the extremes of what is possible, so that few genotypes produce these values, segregation variance will be radically reduced; the derivation of the infinitesimal model depends on there being a very large number of genotypes compatible with each trait value, so that conditioning on the trait does not give significant information about the underlying genotype frequencies.

In order to understand why for ‘typical’ trait values, knowing the trait value for an individual provides very little information about the allelic effect at a particular locus, it is instructive to consider a simple example. The argument we use is similar to that on p.402 of Fisher (1918), where it is expressed in terms of a regression. Suppose that a particular trait is determined by the sum of allelic effects at *M* independent loci, with the allelic effect at the *l*th locus being 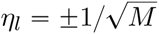 with equal probability. Now suppose that we condition on the trait value being 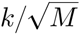. For definiteness, we take *M* and *k* both to be even. What is the conditional probability that 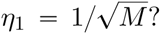 An application of Bayes’ rule gives

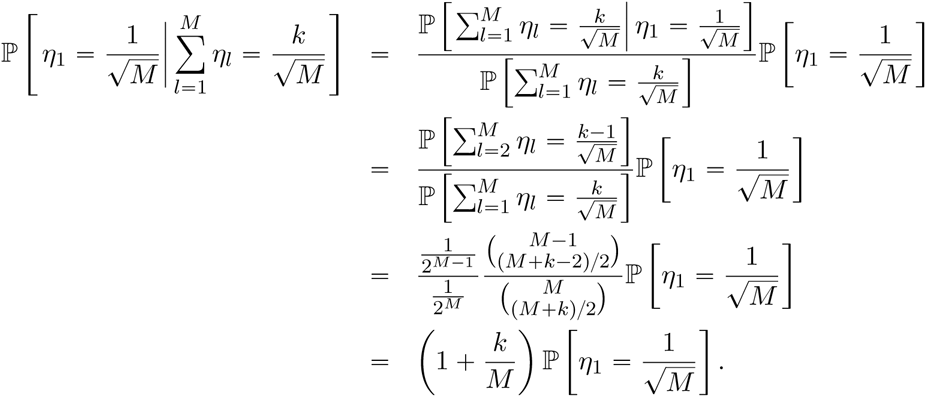

For large *M*, a ‘typical’ value for *k* is 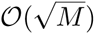, and then this calculation says that, for any particular locus, the chance that it ‘notices’ the conditioning is 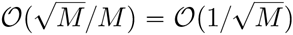. On the other hand, at the extremes of what is possible *(k = ±M*) the value of the trait gives complete information about the allelic effect at each locus.

As we see below, essentially the same argument applies to much more general models for the allelic effects at each locus.

### 3.1 The additive case with mutation and environmental noise

Once we have established our notation, we shall set out the derivation in detail for the strictly additive case but extended to allow (house of cards) mutation. We also suppose that the observed trait value is a combination of a genetically determined component and an environmental noise which, for simplicity, we take to be an independent draw from a mean zero Gaussian distribution for each individual in the population. Laying out this argument carefully enables us to identify the conditions under which our results can be modified to include epistasis. Throughout we concentrate on the haploid case, although (at the expense of more complicated notation and formulae) the same approach extends to the diploid case with dominance.

The formulae that follow are, at first sight, a little daunting. To make them slightly easier to navigate, we impose some conventions in our notation. Table 1 summarises all our notation.

**Table 1.**
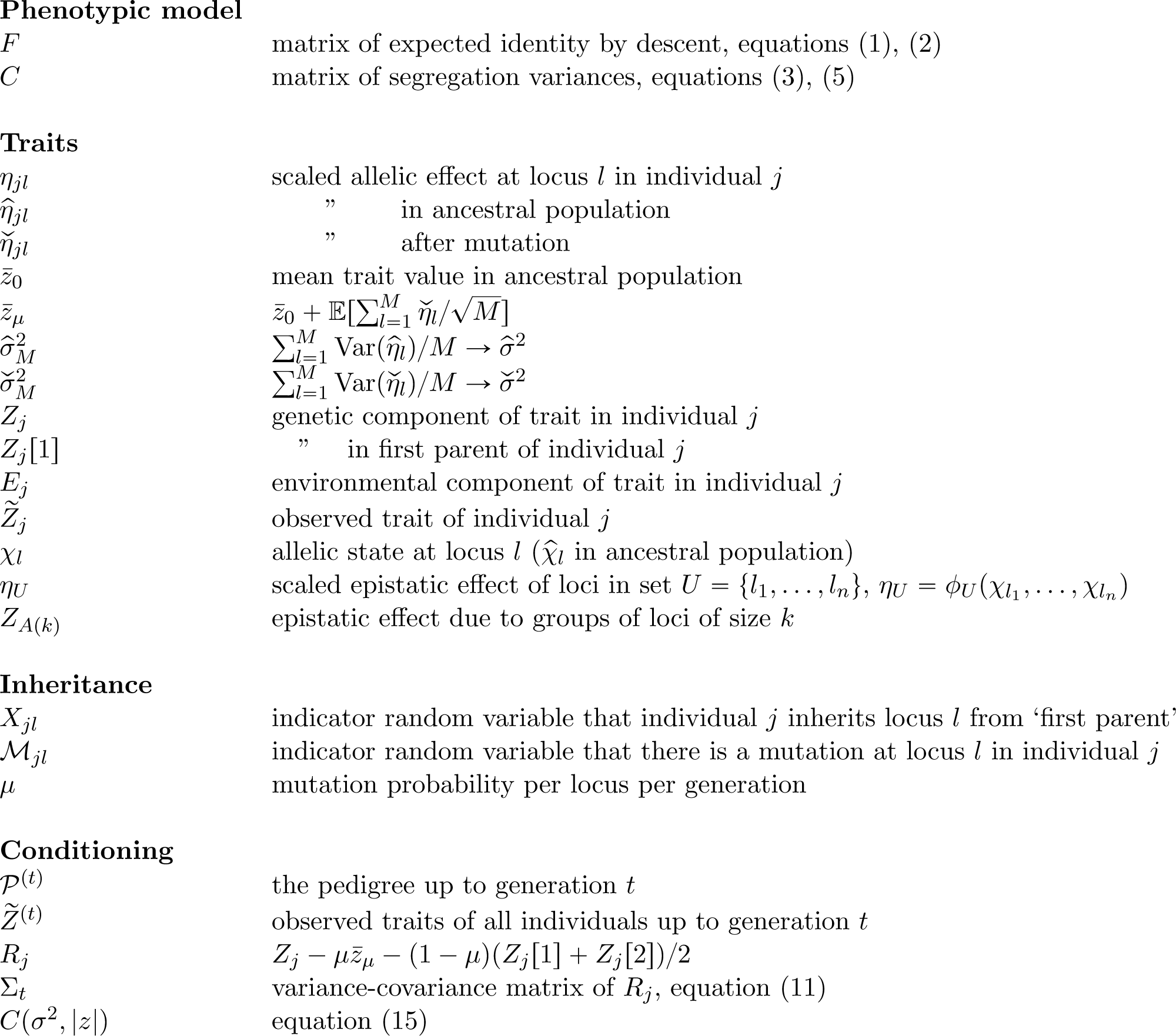

#### Assumptions and Notation

We reserve the indices *i* and *j* for individuals in our population, whereas *l* and *m* are used for loci, of which there are *M*. Generation number will be indexed by *t* (but will mostly be implicit). The total population size in generation *t* is *N*_*t*_.

1. *Allelic effect at locus l*. We denote the allelic effect at locus *l* in the *j*th individual by 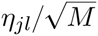. We centre *η*_*jl*_ relative to the mean allelic effect at locus *l* in the ancestral population. The scaling of 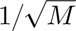 ensures that the additive genetic variance is of order one. The random variable *η*_*jl*_ is assumed to be uniformly bounded over all loci, with |η_*jl*_| ≤ *B*. We sometimes refer to it as the *scaled* allelic effect.
2. *Genetic component of the trait value*. The genetic component of the trait value in the *j*th individual in the present generation will be denoted by *Zj*. It will always be written as 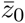, its average value in the ancestral population, plus a sum over loci of allelic effects. That is, in the notation just defined, the genetic component of the trait of the *j*th individual is

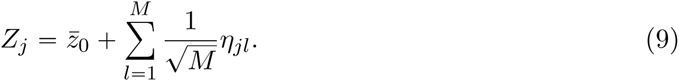
3. *Environmental noise and observed trait value*. We suppose that the observed trait value is

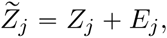

where the *E*_*j*_ are independent normally distributed random variables with mean zero and variance 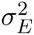.
4. *Ancestral population*. Although it is not strictly necessary, we assume that at time zero, there is an ancestral population of unrelated individuals. The genetic component of the trait value in the *j*th individual in the ancestral population is written as

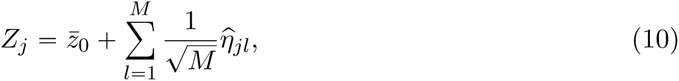

where the 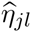 are independent for different values of *j*, with the same distribution as 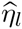 where 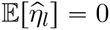 for all *l*. Furthermore, we assume linkage equilibrium in the ancestral population. We shall write

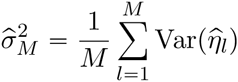

and assume that 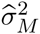 converges to a finite limit 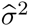 as *M* → ∞.
5. *Parents*. To distinguish the parents of an individual we order them. The symbols [1] and [2] will refer to the first and second parents of an individual, respectively. Thus *η*_*jl*_[1] is the scaled allelic effect at locus *l* in the first parent of the *j*th individual. Similarly, *Z*_*j*_[1] will denote the genetic component of the trait value of the first parent of individual *j*. Note that we allow selfing, in which case parents 1 and 2 are identical.
6. *Mutation*. The mutation probability per locus, per generation, is *μ* (independent of *l*). If there is a mutation at locus *l*, then the scaled allelic effect of the mutant is 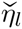. We write 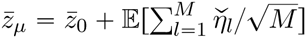 and

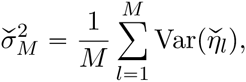

and we suppose that 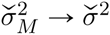 as *M* tends to infinity. If we are referring to a mutation at the *l*th locus in the *j*th individual, we sometimes emphasize this in our notation by writing 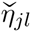.

Centering allelic effects relative to their mean in the ancestral population may seem unnatural, since these are quantities that we could never hope to measure, but in fact it is simply a mathematical convenience: only the variances of these quantities appear in our results.

#### Inheritance

We use Bernoulli random variables to encode the parent from which a locus inherits its allelic type and the presence/absence of a mutation at a particular locus in a given generation. Thus

1. We write *X*_*jl*_ = 1 if the allelic type at locus *l* in the *j*th individual is inherited from the ‘first parent’ of that individual; otherwise it is zero. In particular, ℙ[*X*_*jl*_ = 1] = 1/2 = ℙ[*X*_*jl*_ = 0].
2. We write *M*_*jl*_ = 1 if there is a mutation at locus *l* in individual *j;* otherwise it is zero. Under our assumption of a constant probability of mutation across all loci, we have ℙ[*M*_*jl*_ = 1] = *μ* = 1 - ℙ[*M*_*jl*_ = 0].

Of course there are really independent Bernoulli random variables capturing the inheritance in each generation, but, since we only discuss transitions one generation at a time, we suppress that in our notation.

#### Conditioning

We shall use *P*^(*t*)^ to denote the pedigree relationships between all individuals up to and including generation *t* and 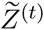 will denote the observed traits of all individuals in the pedigree up to and including the *t*’th generation. We shall be deriving the distribution of trait values in generation *t* conditional on knowing *P*^(*t*)^ and 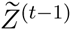 We distinguish *P* from the matrix of identities *F*, because conditional on 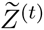, it is no longer true that tracing back through the pedigree, an allele is equally likely to come from either parent; indeed proving that this is *almost* the case is a key part of our derivation.

#### Distribution of the genetic components of trait values

Our first aim is to understand the genetic component of the traits, conditional on *P*^(*t*)^ and 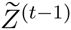. Our precise result will be that conditional on *P*^(*t*)^ and 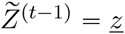,

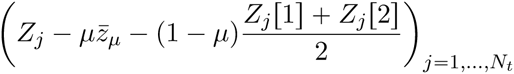

converges (in distribution) to a multivariate normal random variable with mean zero and diagonal covariance matrix Σ_*t*_ with *j*th diagonal entry (Σ_*t*_)_*jj*_ given by the segregation variance among offspring of the parents of individual *j*. We shall present the proof of this result in some detail, starting with the ancestral population and then working recursively through the pedigree. Crucially, we shall keep track of the rate of convergence to multivariate normality at each stage, as it is this that allows us to move from one generation to the next. We remark that since we only condition on 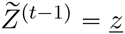, the conditioned quantities *Z*_*j*_[1] and *Z*_*j*_[2] are still random. The expression that we shall obtain for (Σ _*t*_)_*jj*_ is

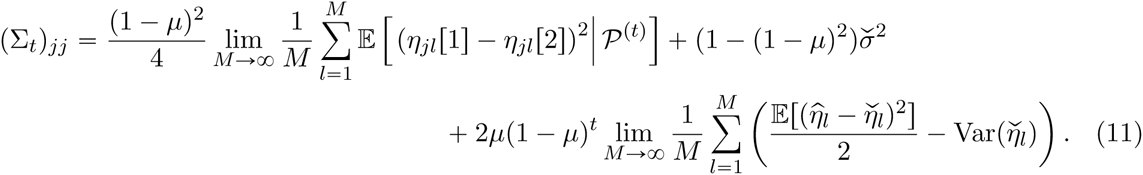

This is precisely the limit as *M* → ∞ of the segregation variance among offspring of the parents of the *jth* individual given by the recursion (5). When *μ* = 0, (Σ_*t*_)_*jj*_ takes the much simpler form

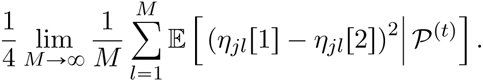

This is just *V*_0_ (1 - *F*_*j*_) where *F*_*j*_ is the probability of identity of the two parents of the *j*th individual and so we recover Equation (3).

#### Genetic component of the trait distribution in the ancestral population

As a first step we show that as the number of loci tends to infinity, the distribution of the genetic component *(Z*_1_,…, *Z*_*N*_0__) of the traits in the ancestral population converges to that of a multivariate normal with mean vector 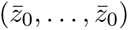 and variance-covariance matrix 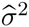 Id, where Id is the identity matrix.

To prove this, it is enough to show that for any choice of *β* =(*β*_1_,…, *β*_*N*_0__) ∈ ℝ^*N*_0_^,

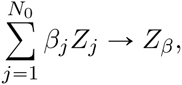

where *Z*_*β*_ is normally distributed with mean 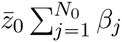 and variance 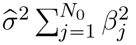. Of course, the result will follow from the Central Limit Theorem, but we need to have some control over the *rate* of this convergence. Moreover, although in the ancestral population the allelic effects 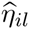 are independent, they do not all have the same distribution, and so we need an extension of the classical Central Limit Theorem. The version that we use is recorded in Appendix A. Not only does it allow non-identically distributed random variables, but it also allows some dependence between them. This is essential in the extension to epistasis. It is convenient to write 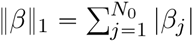 and 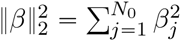. For the ancestral population, Theorem A.2 yields

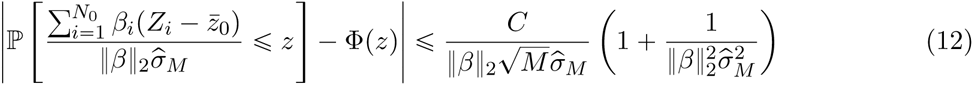

where Φ is the cumulative distribution function of a standard normal random variable and the constant *C* has an explicit expression (depending only on *B* and ‖*β*‖_1_). The details of the proof are in Appendix B.

#### Strategy of the derivation

It will be convenient to define

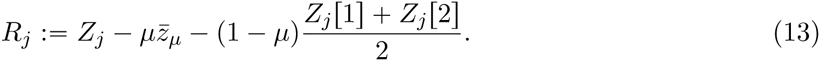

Our proof will be recursive. Suppose that we have our result for generation *(t* - 1). The key step is then to show that conditioning on knowing *P*^(*t*)^ and 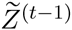 provides negligible information on the values *η*_*jl*_[1], *η*_*jl*_[2] for individual *j* in generation *t*. Through an application of Bayes’ rule, just as in our toy example at the beginning of the section, this will essentially boil down to showing that

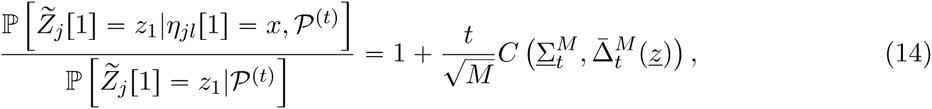

where, since 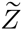 is a continuous random variable, the ratio on the left should be intepreted as a ratio of probability density functions. Here

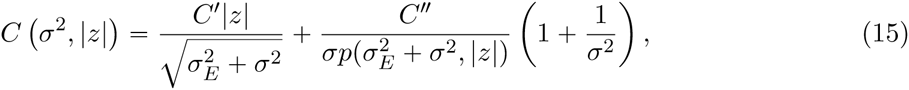

where 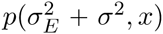 is the density at *x* of a 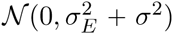 distributed random variable, 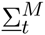 is the minimum segregation variance of any family in the pedigree up to generation *t*, 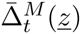 is the maximum over the pedigree up to time *t* of

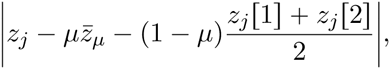

and *C′, C″* are constants, depending only on *B*, our bound on the scaled allelic effects. The proof depends crucially on knowing the rate of convergence of the deviations of the parental trait values to a multivariate normal.

What (14) allows us to deduce (via Bayes’ rule) is that knowing the trait of an individual gives very little information about the allelic state at a single locus. Although intuitively clear, since all loci have a small effect on the trait, this is slightly delicate and indeed, as we see from (15), will break down if the segregation variance somewhere in our pedigree is small or if a trait in the pedigree is too extreme. Armed with (14) we can approximate the distribution of the allelic effects conditioned on *P*^(*t*)^ and 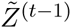 by those conditioned just on *P*^(*t*)^ and then it is an easy matter to identify the limiting variance of the random variables (*R*_*j*_)_*j* = 1,…,*N*_*t*__ in generation *t*.

Convergence of the vector of the genetic components of the trait values in generation *t* to a multivariate normal is then an application of Theorem A.2. Knowing the rate of this convergence allows us to prove (14) for generation (*t* + 1), and so on.

#### One generation of reproduction

We have already proved the asymptotic normality for the ancestral population. To begin the recursion, we consider the first round of mating.

We suppose that, for each *j*, we know the parents of individual *j* and their observed trait values, 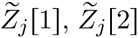. In the notation defined above, this is precisely *P*(1) and 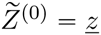. Then we claim that *knowing this information*,

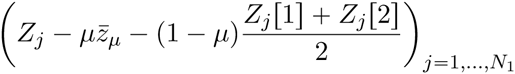

converges in distribution to a mean zero multivariate normal random variable with diagonal variance-covariance matrix Σ_1_ whose on-diagonal entries are given by (Σ_1_)_*jj*_, the segregation variance among offspring of the parents of *j*th individual. By definition, for a given individual *j* in the first generation, we have

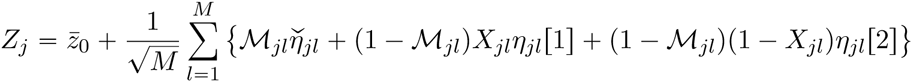

which immediately gives an expression for *R*_*j*_. The random variable *R*_*j*_ satisfies 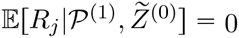. We must calculate its variance. First, we use the normal approximation to the distribution of ancestral traits to show that

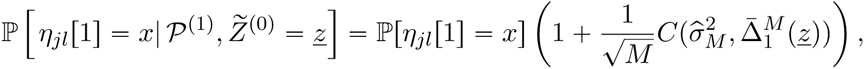

where *C* was defined in (15). Since individuals in the ancestral population are assumed to be unrelated, *η*_*jl*_[1] and *η*_*jl*_[2] are independent (provided the parents are distinct), and combining the calculation above with the symmetric one for *η*_*jl*_[2] we can calculate

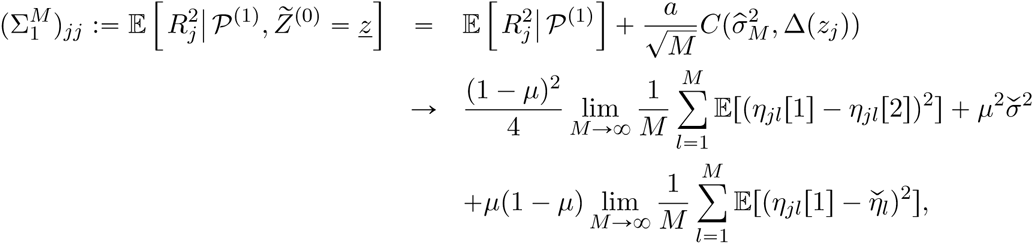

which, since *η*_*jl*_[1] has the same distribution as 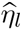, is (11) with *t* = 1. The details are in Appendix D. Since the Bernoulli random variables that describe inheritance in different individuals are independent, it is easy to check that 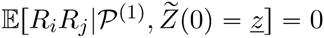.

To verify convergence to a multivariate normal, we mimic what we did in the ancestral population: for an arbitrary vector *β* = (*β*_1_, …, *β*_*N*_1__) we show that 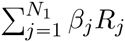 converges to a normal random variable as *M* → ∞. The details are in Appendix D.

#### Generation *t*

We now proceed to the general case. We want to show that conditionally on *P*(*t*) and 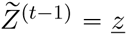, *R*_*j*_ given by (13) converges in distribution as *M* → ∞ to a mean zero, normally distributed random variable with diagonal variance-covariance matrix Σ_*t*_ given by (11). Independence of the Bernoulli random variables that determine inheritance in different individuals once again guarantees that for 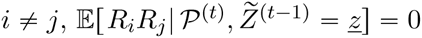. As in generation one, the key is to show that

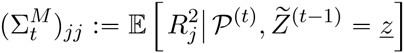

is almost independent of *ẕ*. Convergence to (Σ _*t*_)_*jj*_ given by the expression in (11) is then straightforward (see Appendix E). The only difference from generation one is that in calculating 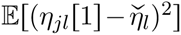 we use that *η*_*j*_[1] is distributed as 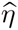 with probability (1 - *μ*)^*t*-1^ and as 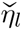 with probability 1-(1-*μ*)^*t*-1^.

The involved step is to estimate

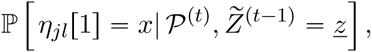

which, again by Bayes’ rule, reduces to checking (14). At first sight, it seems that knowing the trait value of all the pedigree ancestors of the *j*th individual should give us much more information about *η*_*jl*_ than just knowing the parental traits gave us in generation one. In Appendix E we show that this is not really the case. The key is that the *differences* in trait values between individuals that are identical by descent at locus *l* are independent of the scaled allelic effect at locus *l*,sowe don’t accumulate any more information by observing all of these individuals than by observing just one of them.

We can perform entirely analogous calculations for the conditional joint law of *η*_*jl*_[1] and *η*_*jl*_[2]. This enables us to identify the mean and the variance of the limiting distribution of traits in the population and, once again, Theorem A.2 can be used to establish that it is indeed a multivariate normal. In particular,

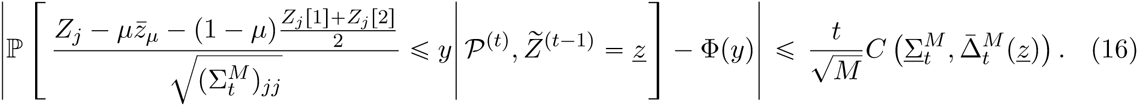

In other words, we have established that the genetic components of the vector of traits follows the infinitesimal model with an error of order 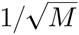 per generation.

Adding an environmental component to the observed traits in the population makes the model more realistic; it also serves a mathematical purpose. As we explain in Remark C.1, without environmental noise, some extra conditions are required to guarantee a rate of convergence of order 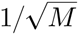 (which is the best possible) to the limiting Gaussian distribution. If they are satisfied, then the calculations that we have just performed are unchanged if we condition on *Z*^(*t*-1)^ the genetic components of the trait values of individuals in the pedigree up to time (*t*-1), instead of the observed values. Our assumption that the environmental noise is Gaussian is certainly unnecessarily restrictive (it would serve the same mathematical purpose if it had any smooth density). It has the advantage that we will be able to obtain explicit formulae for the distribution of observed traits.

#### Observed traits

In the presence of environmental noise, we cannot directly observe the genetic component of the trait. The infinitesimal model as stated above does not apply to the observed trait values. To see why, we write

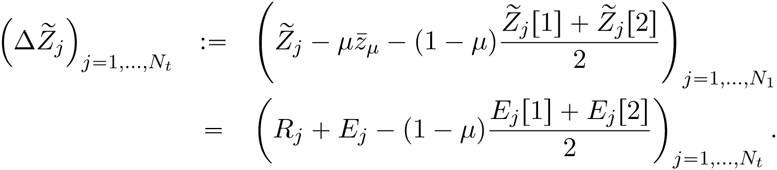

We have already checked that (*R*_*j*_)_*j* = 1,…*Nt*_ is a multivariate Gaussian vector which is (almost) independent of 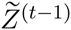, and by assumption the same holds true for (*E*_*j*_)_*j* = 1,…,*N*_*t*__. The difficulty is that the environmental components of the parental traits are *not* independent of the observed traits in generation *(t* - 1). However, under our assumption that the environmental noise is normally distributed, 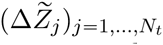 is still asymptotically normally distributed and we can derive recursions for the mean vector and variance-covariance matrices.

In what follows we assume that we are already in the asymptotic regime in which the genetic components of the traits follow a multivariate normal distribution. In fact we are accumulating errors of order 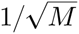 per generation in so doing.

To find the distribution of the observed traits in generation *(t +* 1) conditional on *P*^(*t*+1)^ and 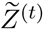, we need to calculate the conditional distribution of the vector (*E*_*j*_)_*j*=1,…,*N*_*t*__. Evidently this vector is independent of 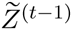 and, granted that we have already calculated the corresponding conditional distributions for the environmental noise in generation (*t*-1), calculating the conditional distribution of (*E*_*j*_)_*j*=1,…,*N*_*t*__ is reduced to applying standard results on conditioning multivariate normal random variables on their sum. In particular, the conditioned vector is still a multivariate normal.

We write 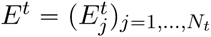 for the random vector

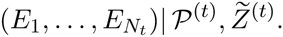

Notice that in contrast to what went before, we are conditioning on knowing all observed trait values up to and including generation *t*. We derive a recursion for the mean vectors *A*_*t*_ and the variance-covariance matrices 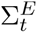 of these conditioned random vectors. Since environmental noise is not transmitted from parent to offspring, this is considerably more straightforward than our previous recursions.

In order to keep track of generations, we suppose that in generation *t* we condition on 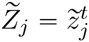. In this notation, the mean of 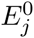 is determined by

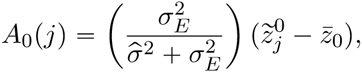

and the variance-covariance matrix of *E*^0^ is

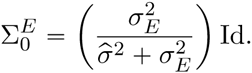

Now suppose that we have calculated *A*_*t*-1_ and 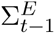. Then

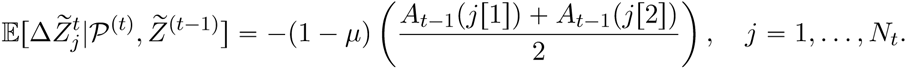

To calculate the variance-covariance matrix, we treat the cases *i = j* and *i* ≠ *j* separately. In the expression below, *i*[*a*] is the *a*th parent of individual *i* (with *a* ∈ {1, 2}). First suppose that *i* ≠ *j*, then

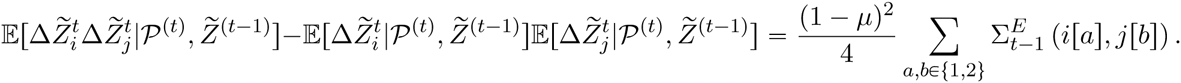

If *i = j*, there are two additional terms corresponding to the variances of *R*_*j*_ and *E*_*j*_,

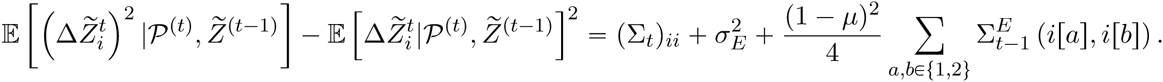

If we denote the variance-covariance matrix of 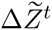 by Σ_*BB*_(*t*), then the recursion which allows us to pass to the next generation reads

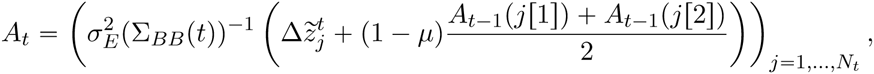

and variance-covariance matrix

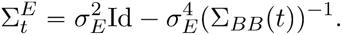

The details of this derivation can be found in Appendix F. Although not as simple as the expressions one obtains for the genetic component of the trait alone, one can now read off the multivariate normal distribution of the observed trait values in generation *t* conditional on *P*^(*t*)^ and 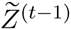. Notice, in particular, that there will be correlations among individuals.

#### Accuracy of the infinitesimal model as an approximation

The infinitesimal model does not just say that the trait distributions in the population can be approximated by a multivariate normal random variable, but it also asserts that the variance of the genetic components of the traits is approximately independent of the trait values of the parents. What our calculations show is that in approximating 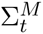 conditional on *P*^(*t*)^ and 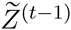 by the same quantity conditioned only on *P*^(*t*)^ (that is the right hand side of (11) before we take the limit), we are making an error of order (recall (14) in particular)

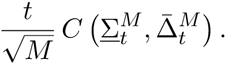

In other words, the infinitesimal model remains valid for order 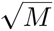 generations, provided that the minimum segregation variance in the pedigree is not too small and none of the traits are too extreme.

### 3.2 Beyond additivity: epistasis

In this section we outline how to extend our arguments to include epistasis. As we noted before this will require some conditions. First, we must avoid *systematic* epistasis, but as long as epistasis is not consistent in direction, we expect that it has negligible effects on the segregation variances. Second, there must be enough independence between loci that we have a Central Limit Theorem: epistatic interactions must be ‘weak’ in an appropriate way.

For simplicity we assume that there is no mutation and no environmental noise. As before, we are going to need control over the rate of convergence of the deviations in trait values to a multivariate normal and for this we shall use Theorem A.2. The conditions of the theorem require that the highest degree of the dependency graph among loci should not be too big. For this reason, we shall suppose that the *M* loci are subdivided into groups of size *D* (independent of *M*) in such a way that two loci from different groups never interact. We write *𝕌*_*g*_, *g* = 1,…*M/D* for these groups. For example, assuming that locus 2*g* - 1 interacts only with locus 2*g* for every *g* ∈ {1,…, *M*/2} corresponds to *𝕌*_*g*_ = {2*g* - 1, 2*g*}, 1 ≤ *g* ≤ *M*/2. In fact, we would expect our result to remain true if we replace this by other ‘sparsity’ conditions, but we are not aware of a convenient form of the Central Limit Theorem without this stronger condition.

We extend our previous notation and for *U* ⊂ 𝕌_*g*_, we write *η*_*U*_ (assumed uniformly bounded) for the scaled epistatic effect of the loci in *U*. We write *χ*_*l*_ for the allelic *state* at locus *l* (of which we assume there are finitely many) and if *U = {l*_1_,…, *l*_*n*_}

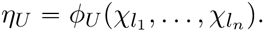

We suppose that in the ancestral population, allelic states at different loci are independent and we shall denote them 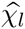. Correspondingly 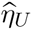 is the scaled epistatic effect of loci in *U* in the ancestral population. In this notation,

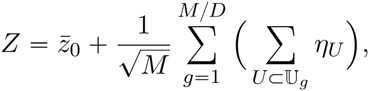

and 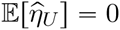 for all *U*.

The other key assumption is the following. Suppose that *U = {l,l*_1_,…, *l*_*m*_}, then for any choice of (*x*_*l*_,…, *x*_*lm*_),

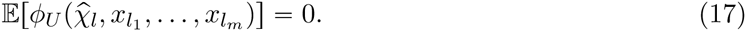

A particular consequence of this is that 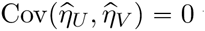 unless *U = V*. Although we use the rather strong symmetry assumption (17) in our proof, we do not believe that it is necessary. Extending our result to more general conditions will be the subject of future work.

Our definition of *∆Z1*_*j*_ (we omit the tilde since there is no noise) must now be extended. It is convenient to group the epistatic components of *Z*_*j*_ according to the number of loci involved. Then

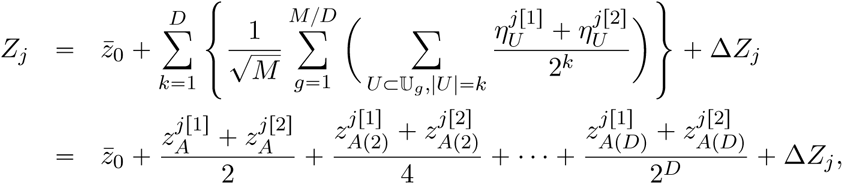

where

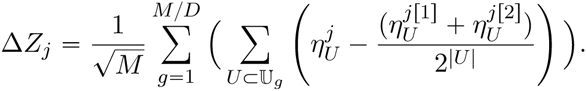

Here *z*_*A*(*k*)_ is the epistatic effect due to groups of loci of size *k*. By changing 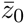 if necessary, we may assume that in the ancestral population the expectation of *z*_*A(k)*_ is zero for all *k*. We assume that we can measure these effects in each parent and our aim is to show that conditionally on knowing 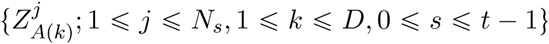, the *D* epistatic effects in every individual trait up to generation *t* - 1, the vector (∆*Z*_1_,…, ∆*Z*_*N*_*t*__) converges to a multivariate normal as *M* → ∞.

The proof follows a familiar pattern. Asymptotic normality of the traits in the ancestral population is immediate from our assumption of independence across loci and Theorem A.2 and the analogue of (14) follows essentially exactly as before.

When we pass to later generations, we encounter two complications. First, it is no longer the case that 𝔼[∆*Z*_*j*_] = 0. Nonetheless, as we explain in more detail in Appendix G, a coupling argument, which exploits the fact that conditional on the *z*_*A*(*k*)_‘s the allelic states have almost the same distribution as the unconditioned one, combined with (17) imply that its expectation is order 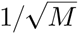.

The second complication is that the ∆*Z*_*j*_ ‘s are no longer asymptotically uncorrelated. In the first generation, correlation comes about if two individuals share the same parents, are identical by descent at all loci from a set *U and* they did not inherit the alleles at all loci in *U* from the same parent. To see this, the chance that they did inherit all the loci from the same parent, *j*[1] say, is 1/2^|*U*|^ and then this term is cancelled by the corresponding centering term, 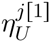. In particular, such correlation requires |*U*| ≫ 2, which is why it did not appear in the additive case. In later generations, correlation again comes from identity at all the loci in *U*, but there may be multiple different routes through the pedigree that result in this, so we no longer require that *i*[1] and *i*[2] are the same as *j*[1] and *j*[2]. However, just as in generation one, the contribution to the correlation will be zero if the individuals share no pedigree ancestry or if one of the individuals inherited all its loci in *U* from the same parent. Thus the contribution from the set *U* takes the form

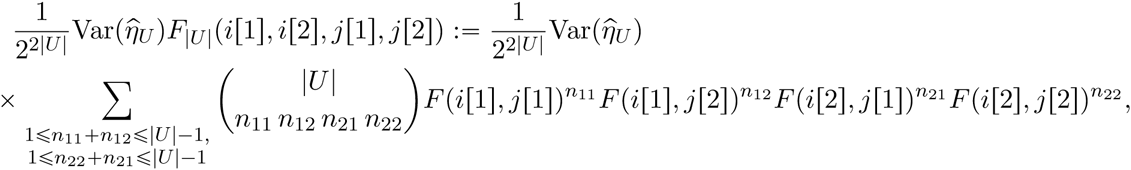

where *F(a, b*) is the probability that individuals *a* and *b* are identical by descent at a given locus and again we have exploited the fact that, knowing the pedigree, the pattern of inheritance is almost unchanged by knowing trait values. Summing over *U*, we obtain that for *i* ≠ *j*, (in the limit as *M* → ∞),

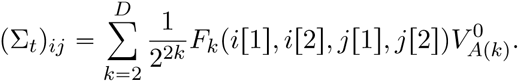

A similar calculation shows that

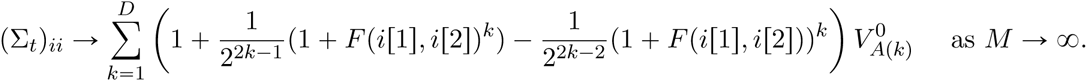

Although notationally complicated, the ideas are the same as in the additive case and we can conclude that conditionally on *Z*^(*t* - 1)^ and *P*^(*t*)^, the vector 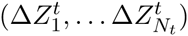 converges in distribution as *M* → ∞ to a multivariate Gaussian with mean zero and variance-covariance matrix Σ_*t*_. Armed with the speed of convergence (which is again order 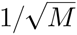) and Bayes’ rule, we can prove that conditioning on *z*_*A*_, *z*_*A*(2)_, … *z*_*A*(*D*)_ provides little information about the allelic state at any particular locus and then use the unconditioned distribution of allelic states to estimate the variance-covariance matrix for the next generation, and so on.

### 3.3 Numerical examples

In this section, we illustrate our results with some numerical examples. We focus on cases with no environmental noise or mutation. We begin with an example that shows how the genetic variance is much less sensitive to selection than the mean. We then show how the genetic variance scales with the number of loci, and finally, show that the variance amongst offspring depends only weakly on the parents’ trait values.

All our examples are based on simulations of a Wright-Fisher model of 100 haploid individuals. With the exception of Figure 3, which investigates how rapidly the infinitesimal limit is approached as the number of loci increases, we set the number of loci *M* = 1000. Our choice of parameters is constrained by computational limits. We mostly simulated equal ‘main’ effects, corresponding to the allelic effects 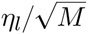 in our derivation of Section 3 taking the values ±α (where 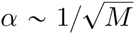), because the infinitesimal limit is then approached with a minimal number of loci. Exponentially distributed effects might be more realistic, but require an order of magnitude more loci to approach the infinitesimal limit (see Figure 3).

We also consider pairwise epistasis. We expect the infinitesimal model to be an accurate approximation if the epistatic effects are sufficiently sparse. Our example is constructed in two stages. First, independently for each ordered pair (*l, m*) of loci with *l* ≠ *m*, with probability 1/*M* we declare it to have a non-zero epistatic effect. Then, each non-zero interaction is assigned an independent sample from a normal distribution with mean zero and standard deviation 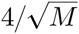. With this construction *γ*_*lm*_ ≠ *γ*_*ml*_. The trait is now defined as *z = δ*·*α + δ*·*γ*·*δ*^*T*^, where the entries in the vector *δ*, which records the genotype, are ±1/2. The epistatic and main effects were scaled with respect to the number of loci, *M*, so that both the additive and epistatic variances are of order 1. This model will not quite satisfy the conditions of our theoretical result, but instead we use it to support our claim that other forms of ‘sparsity’ should be sufficient for the infinitesimal model to provide a good approximation.

We chose *N* = 100 haploid individuals, in order to be able to follow the matrix of identities. Since we have no mutation, variation then dissipates over about *N =* 100 generations. We ran simulations either with no selection, or with directional selection *β* = 0.2, so that fitness *W* is proportional to *e*^*βz*^. Simulations were started with allele frequencies drawn from a Beta distribution with mean *p* = 0.2 and variance 0.2*pq*.

With selection on a heritable trait, fitness is also heritable, which speeds up the loss of genetic variance due to random drift (Robertson, 1961); the variance declines in proportion to 1/*N*_*e*_ per generation, where *N*_*e*_ is somewhat less than the census number. However, in our examples, the variance in fitness, *β*^2^*V*_*G*_, is small, and so the relatedness, *F*, is not appreciably increased by selection.

Figure 1 shows how selection affects the mean and the components of the variance of a trait that is determined by *M* = 1000 loci, in a population of *N* = 100 haploid individuals. Selection on an additive trait changes the mean by 4.5 genetic standard deviations, 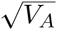, over 100 generations whilst the additive genetic variance decreases by a factor 0.355 (top row). Crucially, this is almost the same as the decrease with no selection, 0.364, and close to the neutral expectation (1 - 1/*N*)^100^ = 0.366. The bottom row shows an example with sparse pairwise epistasis. The additive variance is much higher, and the mean now changes by 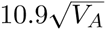 over 100 generations. In both the additive and epistatic examples, the change in mean per generation is close to the predicted 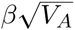. With epistasis, but no selection, the additive variance decreases only by a factor 0.548 over 100 generations (upper dashed line at bottom right), because non-additive variance is ‘converted’ into additive variance. Now, selection does substantially reduce the additive variance, after about 30 generations.

**Figure 1:**
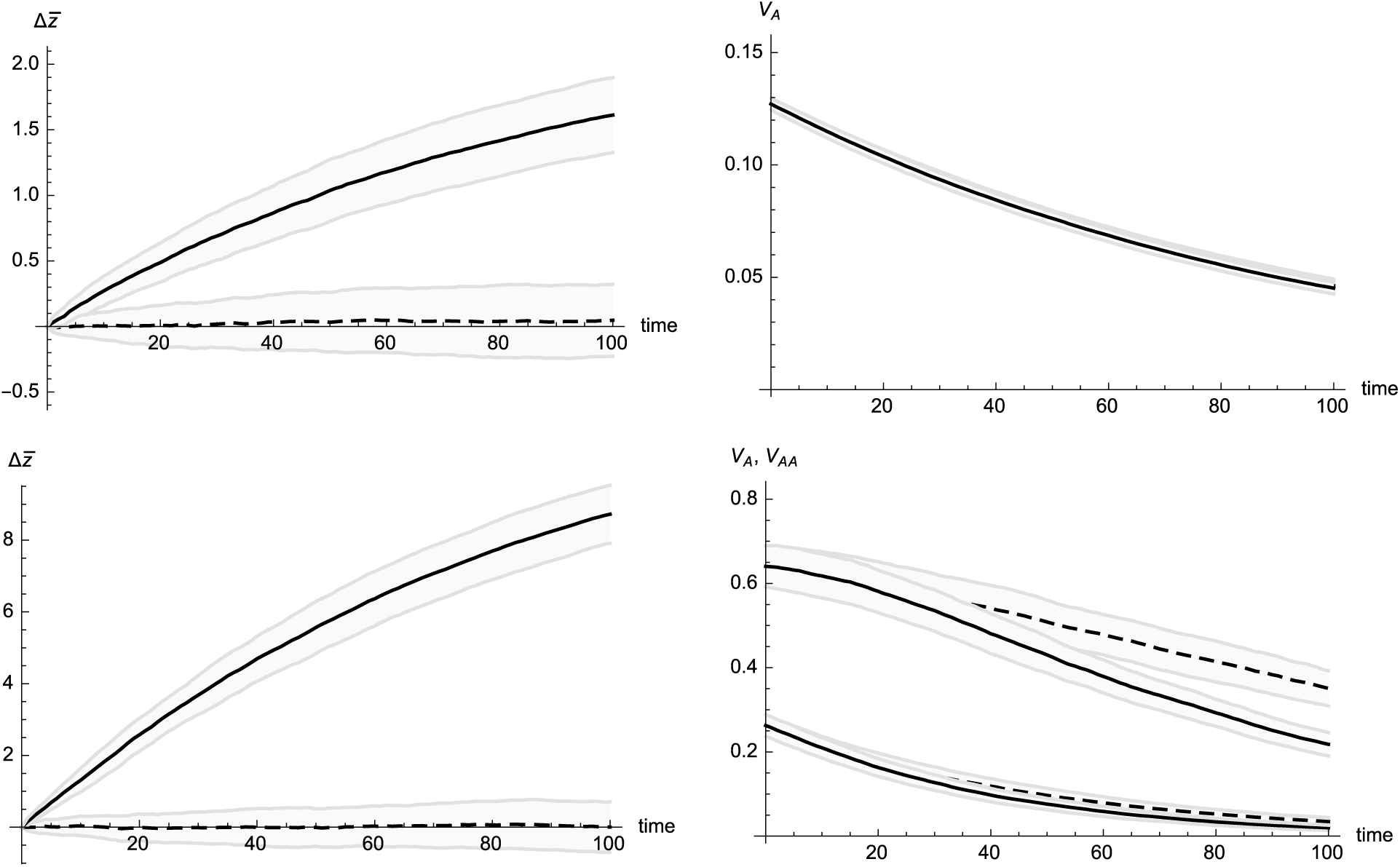
The effect of selection on the mean and variance components. Each panel contrasts directional selection, *β* = 0.2, on a trait (solid line) with the neutral case (dashed line); shaded areas indicate ±1 standard deviation. The top row shows the additive case, with equal effects but random sign, 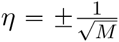; in this example, *M* = 1000 loci, *N =* 100 haploid individuals. The lower row shows an example of sparse pairwise epistasis, as described in the text. The left column shows the change in mean from its initial value, and the right column shows the additive variance, *V*^*A*^, and with epistasis, the additive × additive variance, *V*_*AA*_ (lower pair of curves at bottom right). Only the genic components of variance are shown; random linkage disequilibria produce substantial fluctuations, but make no difference on average (results not shown). Initial allele frequencies are drawn from a U-shaped Beta distribution with mean *p* = 0.2 and variance 0.2*pq*. Individuals are produced by Wright-Fisher sampling, from parents chosen with probability proportional to *W* = *e*^*βz*^. For each example, ten sets of allelic and epistatic effects are drawn and, for each of those, ten populations are evolved; this gives 100 replicates in all.

A cornerstone of our derivation of the infinitesimal model was the result that the distribution of allele frequencies is hardly affected by selection. Figure 2 shows the change in allele frequency over time. In the presence of epistasis, the marginal effect of an allele on the trait depends on current allele frequencies at other loci. In particular, alleles that have a positive effect on the trait in the initial population may have a different effect in the final generation. In order to determine whether the allele frequency spectrum is substantially biased towards alleles that have a positive overall effect on the trait, in the presence of epistasis we recalculate the marginal effects of each allele at the final generation and relabel them accordingly. Over 100 generations, random drift fixes alleles that are rare or common, leaving a flat distribution of frequencies of segregating alleles (grey lines at middle and bottom). Selection shifts the distribution in favour of alleles with a positive effect on the trait. In the additive case, the shift is symmetrical, so that there is virtually no change in the additive genetic variance: *𝔼*[*pq*] stays the same. In contrast, selection on an epistatic trait reduces the overall frequency of segregating alleles. This may be because with epistasis, allelic effects vary with allele frequency, so that alleles experience an additional random fluctuation that reduces diversity.

**Figure 2:**
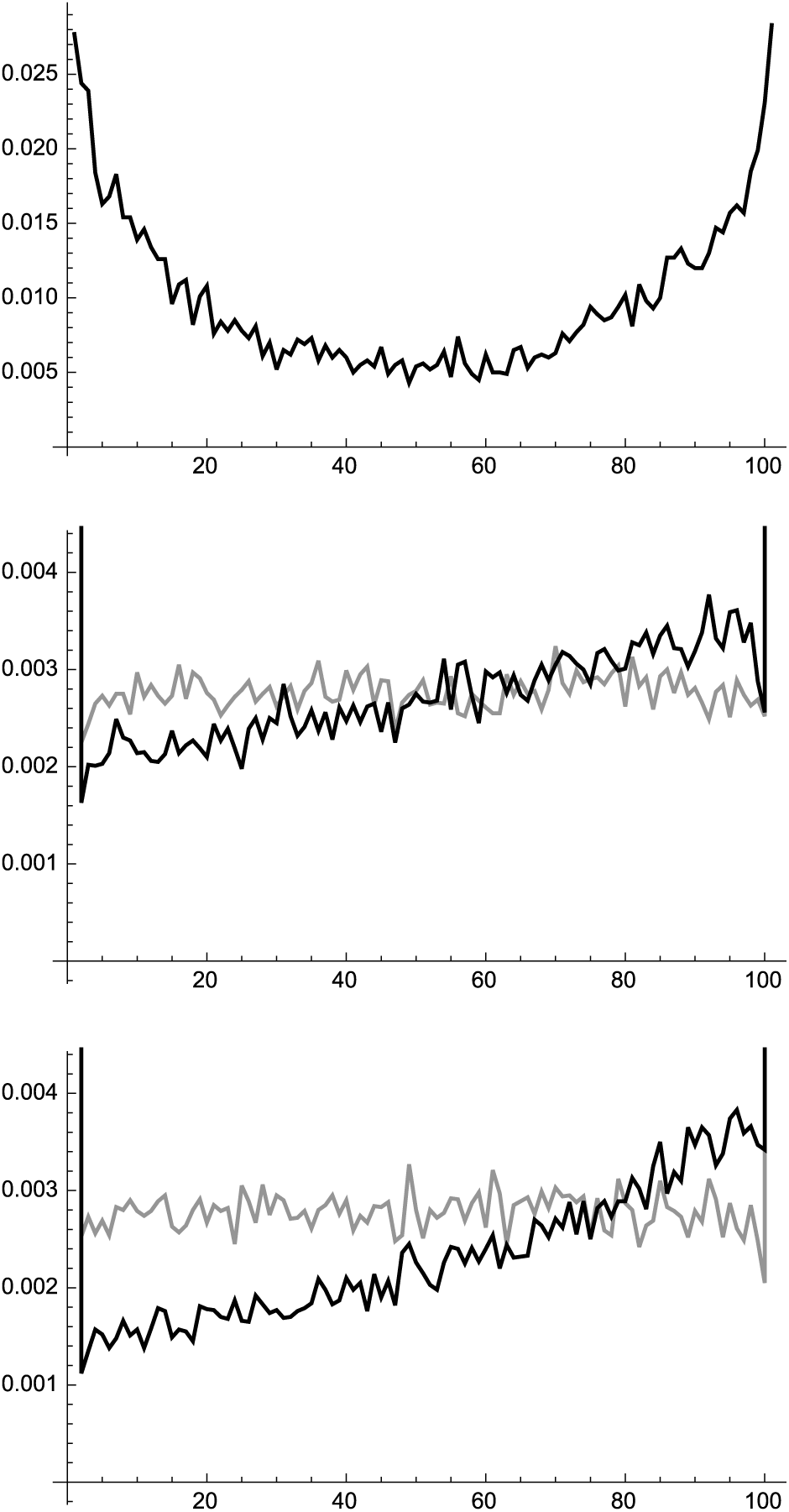
Allele frequency distributions for the examples in Fig. 1. The horizontal axis shows the number of copies of the + allele (0,…, *N*) in a population of size *N =* 100. The vertical axis is the proportion of the *M* = 1000 loci with that number of + alleles. Top: Initial allele frequency distribution, which is independent of allelic effect. Middle: Additive case, after 100 generations, with no selection (grey) or with selection *β* = 0.2 (black). Bottom: With epistasis, after 100 generations where an allele is defined to be + if its marginal effect in the final generation is positive (see main text).

With no selection, the rate of decrease of variance, -∂_*t*_log(*V*_*A*_), is close to 1/*N*, independently of the number of loci (compare small dots with dashed line in Figure 3). With selection on a small number of loci (*M* = 30; left of Figure 3), the additive variance declines much faster, as favourable alleles are swept to fixation. The excess rate of decrease is inversely proportional to the number of loci: 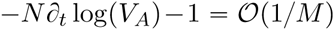; the exponent of the relation, estimated by regression on a log-log scale, is −1.007, −1.019, −0.965 for the three examples of equal effects, exponentially distributed effects, and equal effects plus epistasis. Note that as in Figure 1, the additive variance is much more sensitive to selection in the presence of epistasis (compare upper large dots with medium black dots). However, in both cases the additive variance scales as close to 1/*M*.

**Figure 3:**
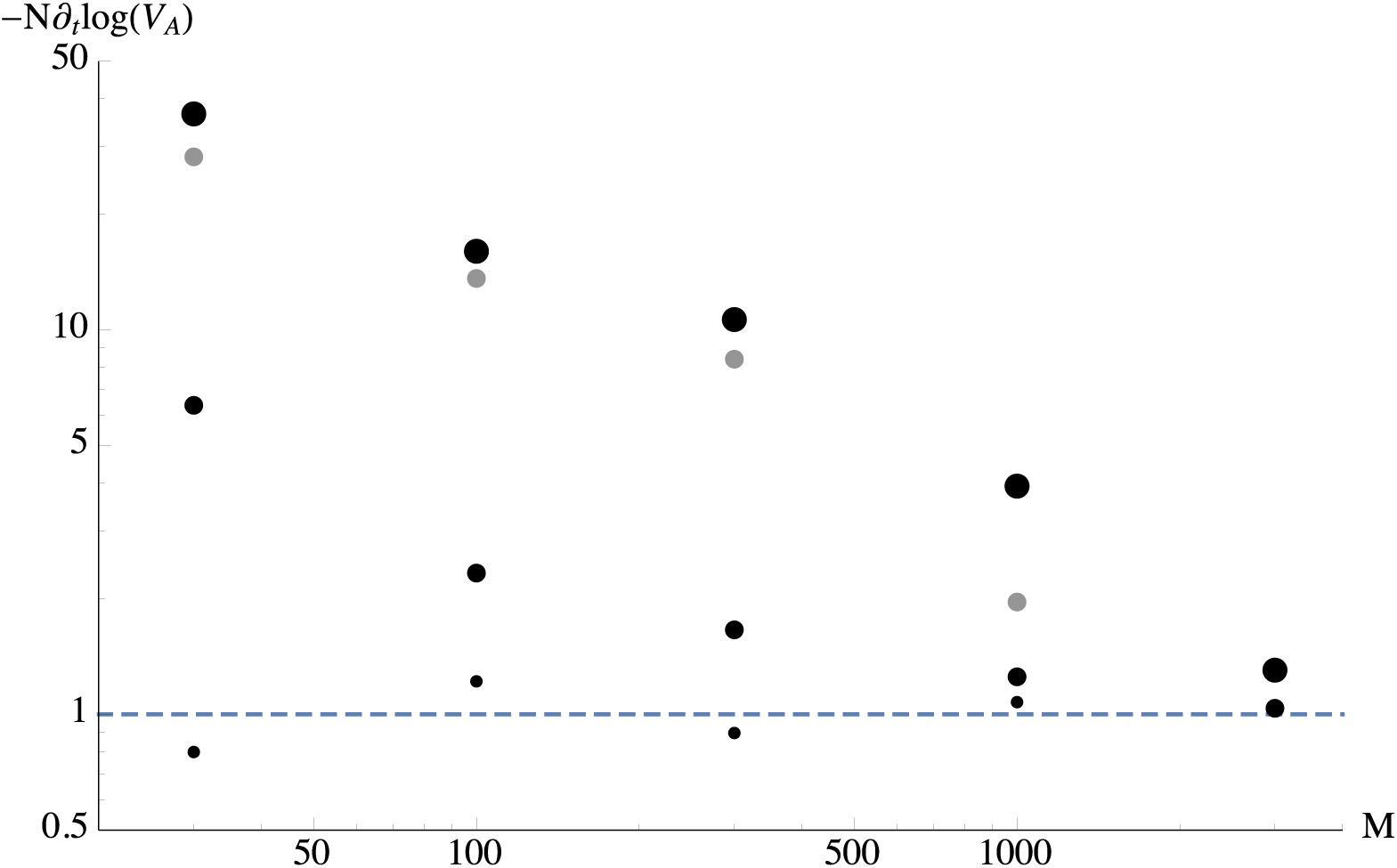
The effect of selection on the genetic variance decreases with the number of loci, *M*. The rate of decrease of additive genetic variance, multiplied by population size, *N*∂_*t*_log(*V*_*A*_), is plotted against *M* on a log-log scale. The neutral rate of decrease of variance is indicated by the dashed line at *N*∂_*t*_log(*V*_*A*_) = 1. Small dots show the neutral case, with equal and additive effects and random sign, 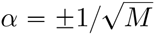. Medium black dots show directional selection *β* = 0.2, again with equal and additive effects, 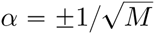. Medium grey dots show *β* = 0.2, with additive effects drawn from an exponential distribution with mean 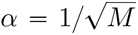 and random sign. Large dots show *β* = 0.2, with equal allelic effects 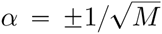, plus sparse pairwise epistasis, as described in the text. Initial allele frequencies are drawn as in Figure 1. The rate of decrease of variance is estimated by regression over 100 generations, *V*_*A*_ being averaged over 10 replicates. These replicates are evolved independently, starting from the same randomly chosen allelic effects, *α, γ*, and initial allele frequencies. Only the genic component of the additive variance is shown.

This is a much faster approach to the infinitesimal model than the upper bound of 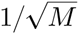 set by our mathematical results. By considering the argument at the beginning of Section 3 for the additive model with equal main effects, we can see that the rate 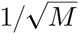 cannot be improved upon in our general result (which applies even when we consider individual families and condition on ancestral traits). However, when, as in Figure 3, we consider the variance across the whole population, we can expect faster convergence. We can understand why this is as follows. The genic component of the additive variance is 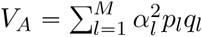. Since the rate of change of the allele frequency at locus *l* due to directional selection *β* on the trait is *βα*_*l*_*p*_*l*_*q*_*l*_, we have 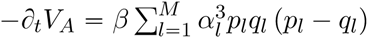 (ignoring the change in the marginal allelic effect due to epistasis). We assumed that *α*_*l*_ has random sign, so that the expected initial rate of change is zero; indeed, when we sample replicates with different effects, *α*, the rate of decrease measured over the first 10 generations is closely correlated with 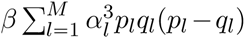, and fluctuates in sign (results for *M* = 30 not shown); the standard deviation of this initial rate scales as 1/*M*. However, the rates of decrease measured over 100 generations are almost all positive, and vary much less than does the initial prediction 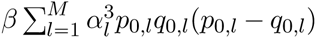, based on the initial allele frequencies. This consistently positive rate of decrease arises because the allele frequencies *p*_*l*_ become correlated with the allelic effect, *α*_*l*_: with no mutation, favourable alleles eventually become common, so that the additive variance decreases as they move to fixation.

For the rest of our examples, we simulated a population of *N* = 100 haploid individuals with *M* = 1000 loci. The main allelic effects at each locus are taken to be 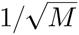, and examples with sparse epistatic interactions are constructed exactly as above. Simulations were run for 100 generations, tracking the full pedigree. We then calculated the matrix of pairwise identity by descent, *F*, relative to the initial population, and also relative to generation 80. We assume that the probability that two pairs of genes at different loci are both identical by descent is *F^2^*.

In Figure 4, we plot the mean and variance within families of the additive and non-additive components against the average of the corresponding components in the two parents; parameters are as in the epistatic example shown in Figure 1. We choose pairs of unrelated parents so that under the infinitesimal model the variances should be constant across pairs. Even in the presence of epistasis, the mean of the additive component in the offspring must be precisely the mean of the additive component in the parents (top left). The mean additive × additive component in the offspring equals half of the mid-parental value of that component, plus a random component that varies between pairs of parents. The slope of the regression line in the top right panel is 0.58, reasonably close to the expectation. The variance of the additive and non-additive components amongst offspring is independent of the mean parental components (bottom row).

**Figure 4:**
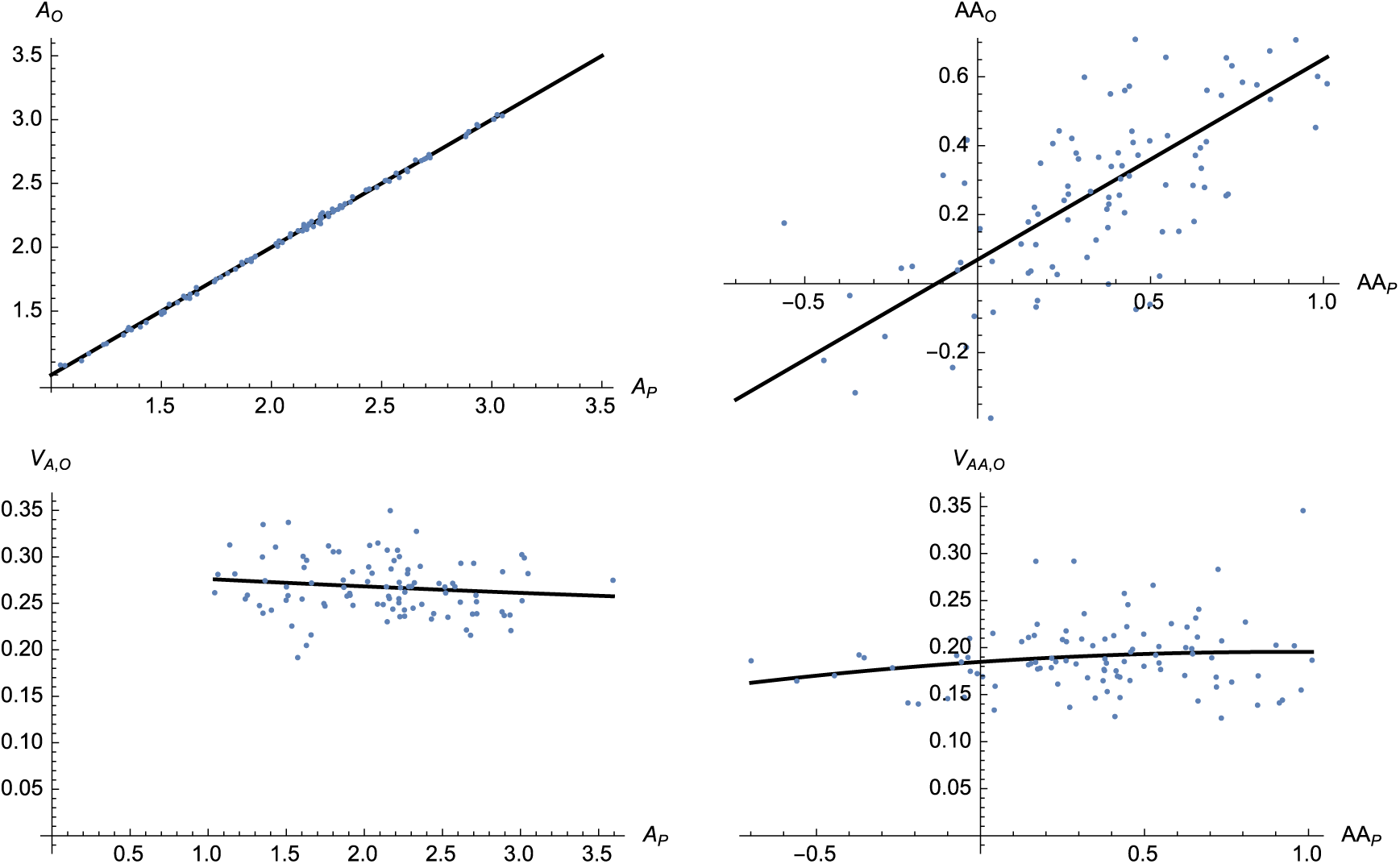
The mean and variance of offspring traits, plotted against components of the parents’ trait values. Top left: Additive component of offspring, *A*^*O*^, against the mean of the parents’ additive component, *A*_*P*_. The line represents *A*_*O*_ = *A*_*P*_. Top right: The same, but for the additive × additive components. The line shows a linear regression. Bottom left: Additive variance amongst offspring, *V*_*A, O*_ against the mean additive components of the parents, *A*_*P*_. Bottom right: Additive × additive variance of offspring against the mean additive × additive component of the parents. Lines in the bottom row show quadratic regressions. The example shows a non-additive trait under selection *β* = 0.2, with *M* = 1000 loci and *N* = 100 haploid individuals, as in Figure 1 (bottom row). At generation 20, 100 pairs of minimally related parents *(F* = 0.16) were chosen, and 1000 offspring were generated for each pair. For each offspring, the components of trait value were calculated relative to the allele frequencies, *p* in the base population. Defining genotype by *X* ∈ {0,1}, these components are *A* = *ζ*·(*α* + (*γ* + *γ*^*T*^)·(*p* - ½)), *AA* = *ζ·γ·ζ*^*T*^, where *ζ* = *X - p*, and the allelic effects *α, γ* are drawn as for Figure 1.

Finally, in Figure 5, we plot the decline of the additive and non-additive variance with relatedness, *F*. Under the infinitesimal model, the within-family additive variance is 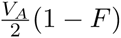, where *F* is the probability of identity between the genomes that unite at meiosis. With sparse pairwise epistasis, the within-family non-additive variance is 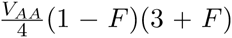 (this expression can be obtained by the same analysis as in Section 3.2, where we compute the non-additive variance due to groups of genes inherited from different parents). Figure 5 is based on a population of 100, simulated for 20 generations under selection *β* = 0.2, as for Figure 4; the average relatedness is then *F* = 0.18 ≈ 1 - (1 - 1/*N*)^20^. The theoretical predictions shown by the solid curves are based on the additive and non-additive variance components in the base population. There is a small deviation from these predictions, because the observed genetic variances include random linkage disequilibria amongst the 100 sampled individuals, whereas the predictions are based on the allele frequencies in the base population.

**Figure 5:**
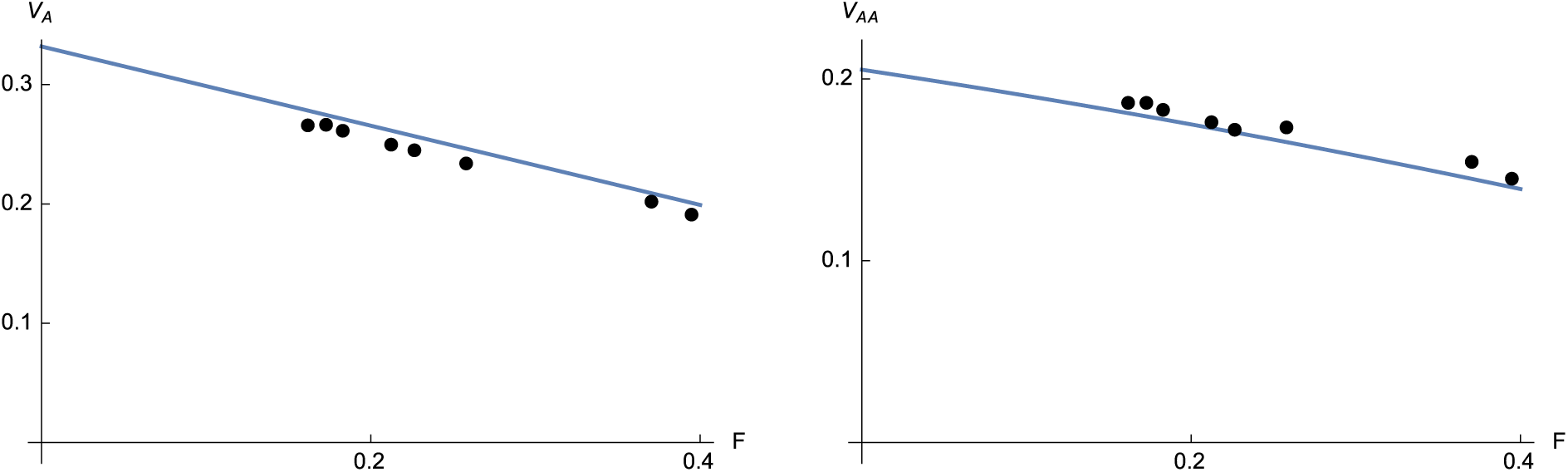
Additive and non-additive variance amongst offspring declines with pairwise relatedness between parents. Solid lines show the theoretical expectations: 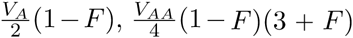. Values are based on a population of 100 individuals, generated as in Figure 4. The 4950 distinct pairs of parents are pooled according to their relatedness, and the variance amongst offspring is estimated from 1000 offspring produced by each pair.

## 4 Discussion

Typically, the distribution of a quantitative trait amongst offspring is Gaussian, with a mean intermediate between the parents’ trait values, and a variance that is independent of those values. This observation goes back to Galton (1877), and was explained by Fisher (1918) as being the result of a large number of unlinked loci with additive effects. The variance amongst offspring depends on the relatedness of the parents, which can be predicted from the pedigree. This infinitesimal model thus provides a complete description of the short-term evolution of quantitative traits, which is independent of any knowledge of the genetics.

We have set out a rigorous justification for the infinitesimal model, which clarifies the conditions under which it holds. These are very general. In the additive case, we can include arbitrary selection and population structure, provided that the segregation variance is not too small and traits are not too extreme. The derivation includes mutation and environmental noise. Most important, the argument that the distribution of the trait amongst offspring is insensitive to selection carries over to allow epistasis. With epistasis, we must now specify a set of variance components, which predict the variance amongst offspring on an arbitrary pedigree. In all cases, the mathematical analysis shows that the infinitesimal model holds up to an error which is at most of the order of 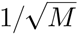, where *M* is the number of loci, while Figure 3 suggests that in some cases the error could be as small as 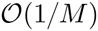.

We have not considered dominance here. With dominance, the variance components no longer suffice to predict the offspring distribution: more complex quantities are involved (Barton and Turelli, 2004). Nonetheless, the proof of convergence of trait values on a given pedigree to a multivariate normal given by Lange (1978) does include dominance and we anticipate that our central result still holds in the limit of a large number of loci: the offspring distribution is independent of the parents’ trait values, and hence insensitive to selection. This, along with a more thorough mathematical investigation of the most general conditions under which epistatic interactions do not disrupt the infinitesimal model, will be the subject of future work.

We have assumed throughout that all loci are unlinked. This is of course inconsistent with assuming a very large number of loci on a finite genetic map. Linkage will reduce the rate at which segregation variance will be released, and more seriously, the breeding value and within-family variance are no longer sufficient to predict the evolution of the population: associations will build up between linked loci, and will only slowly be dissipated. However, in the limit of a long genetic map, selection still has negligible effect on the genetic variance (Bulmer, 1974; Santiago, 1998).

One can imagine an extension of the infinitesimal model to linear genomes, which can readily be implemented in an individual-based simulation. Imagine two long genomes of map length *R*, which initially have a certain total effect on the trait, *z*_1_, *R*]), *z*_2_([0, *R*]). Recombination between these genomes generates a gamete that is a mosaic of blocks derived from one or other parent, {*z*_1_([0, *x*_1_]), *z*_2_([*x*_1_, *x*_2_]),…}. Conditional on the breakpoints, the values associated with the segments *z*_*i*_([*x*_*i*_, *x*_*i*+1_]), form a multivariate Gaussian, conditioned on the sum *z*_*i*_([0,*R*]). At the level of the population, this model is implicit in studies of the effects of background selection, in which heritable fitness variance due to deleterious mutations is spread over the genome (e.g. Good and Desai, 2014).

The infinitesimal model requires that a sufficient number of loci contribute to the trait. With strong inbreeding, the number of contributing loci may become too small for the model to be accurate. This may be a particular problem if variance is contributed by rare recessive alleles: only a few such alleles may contribute in a cross between two close relatives. Thus, while the infinitesimal model may remain accurate for the population as a whole, it might break down under strong inbreeding between particular individuals or in particular subpopulations.

For how long can we expect the infinitesimal model to be accurate? The wide use of the model in animal breeding suggests that it is accurate (or at least, useful) for many tens of generations. Indeed, the sustained response to artificial selection that is typically seen is the strongest support for the infinitesimal approximation (Barton and Keightley, 2002). Remarkably, Weber and Diggins (1990, Fig. 4) found that for a wide range of traits and model organisms, the response to selection over 50 generations is close to the infinitesimal prediction. Responses tend to be slightly below the prediction, suggesting that selection is reducing the variance faster than expected by random drift, but the closeness of the fit implies that most of the selection response is due to alleles that are influenced mainly by random drift (i.e., that *N*_*e*_*s* < 1 or less).

This evidence comes from relatively small populations, and short timescales. In the longer term, mutation becomes significant, and the infinitesimal model predicts a genetic variance in a balance between mutation and drift of *N*_*e*_*V*_*m*_ for a haploid population. This cannot plausibly explain observed heritabilities in large natural populations, since genetic variances do not show a strong increase with population size. (Though, we note that sequence diversity also shows a weaker increase with census population size than expected from naive neutral theory. It is not clear whether quantitative genetic variance increases in proportion with sequence diversity; Frankham, 1996; Willi et al., 2006). It is widely believed that genetic variance is due to a balance between mutation and selection against deleterious mutations. However, it is not clear whether selection acts on the trait or on the pleiotropic effects of the alleles involved, and the contribution of balancing selection of various kinds is unknown (Johnson and Barton, 2005). The infinitesimal model may remain accurate at least for times shorter than 1/*s*; however, the effects of selection at the underlying loci need further theoretical investigation. Estimates of the distribution of fitness effects (largely based on evidence from Drosophila) suggest that there may be a significant fraction of very weakly selected alleles (e.g. Loewe and Charlesworth, 2006); if these contribute to traits as well as to fitness, then the infinitesimal model may hold for long times. However, Charlesworth (2015) concluded that the quantitative genetics of fitness variation in Drosophila can only be reconciled with estimates of fitness effects from population genomics if most fitness variance is either due to relatively strongly selected mutations (*N*_*e*_*s* ≫ 1), or to the side-effects of balancing selection.

The enormous efforts put into mapping quantitative trait loci (QTL), and more recently, to finding associations between genome-wide markers and quantiative traits (GWAS), have identified many QTL, but typically, have not explained much of the genetic variance. There is no mystery about this “missing heritability”: it is to be expected if genetic variance is due to large numbers of alleles of small effect (Yang et al., 2010). Thus, it may be impossible to identify most of the individual alleles responsible for trait variation, even if the whole genome and the whole population is sequenced. Nevertheless, a regression of trait on sequence can significiantly improve predictions of breeding value, even when individual loci cannot be identified: this is the basis of “genomic selection” (Meuwissen et al., 2013). It may be that natural selection is in just the same position as a breeder: selection may change the mean rapidly and predictably, even when the probability distribution of any particular allele frequency is hardly perturbed.

## Acknowledgement.

The authors thank Bill Hill for valuable comments on a previous draft of the paper. A.V. would like to thank Olivier Martin and the members of a working group on the infinitesimal model (in particular Patrick Phillips, Gaël Raoul and Henrique Teotonio) for interesting discussions on this topic.

## A A Central Limit Theorem

The version of the Central Limit Theorem that we use is from Rinott (1994). At the expense of stronger conditions (which for us are fulfilled as a result of putting a uniform bound on scaled allelic effects), this result improves the rate of convergence in the corresponding result in Baldi & Rinott (1989). In contrast to the classical result, it allows for sums of non-identically distributed random variables with some local dependence structure which is most conveniently expressed through what is known as a *dependency graph*.

**Definition A.1** *Let {X*_*l*_;*l ∈ V} be a collection of random variables. The graph 𝒢 = (V, ε), where V and ε denote the vertex set and edge set respectively, is said to be a* dependency graph *for the collection if for any pair of disjoint subsets A*_1_ *and A*_2_ *of V such that no edge in ε has one endpoint in A*_1_ *and the other in A*_2_, the sets of random variables {*X*_*l*_;*l ∈ A*_1_} *and {X*_*l*_; *l ∈ A*_2_} *are independent*.

The degree of a vertex in the graph is the number of edges connected to it and the maximal degree of the graph is just the maximum of the degrees of the vertices in it.

**Theorem A.2 (Theorem 2.2, Rinott (1994))** *Let Y*_1_,…,*Y*_*M*_ *be random variables having a dependency graph whose maximal degree is strictly less than D, satisfying 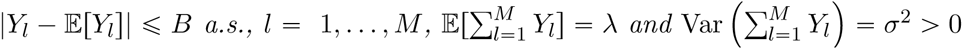. Then*

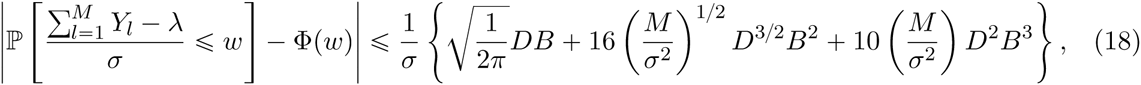

*where Φ is the distribution function of a standard normal random variable*.

In particular, when *D* and *B* are order one and *σ*^2^ is of order *M*, as is the case in our applications, this yields a bound of order 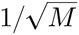.

Although much of the appeal of this result is that it allows dependence between the variables, we shall also use it in the independent case. In that setting, the dependency graph has no edges, and so the maximum degree of any vertex is zero and we may take *D = 1*.

**B Trait distribution in the ancestral population**

In this section, we show that as the number of loci tends to infinity, the distribution of the traits *(Z*_1_,…, *Z*_*N*_0__) in the ancestral population converges to that of a multivariate normal with mean vector 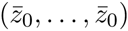 and covariance matrix 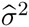 *Id*. To this end, take *β* = (*β*_1_,…, *β*_*N*0_) ∈ ℝ^*N*^0^^ and recall our notation 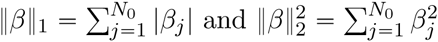. We consider 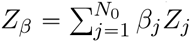.

To apply Theorem A.2, we must first identify the mean and the variance of *Z*_*β*_. Since we are considering the ancestral population, we have

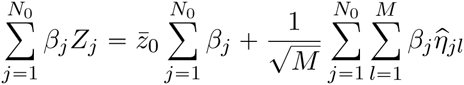

The double sum has mean zero and, since the summands are all independent, variance

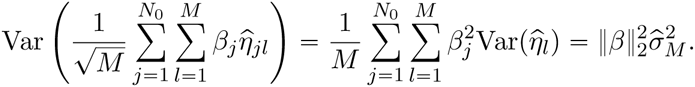

We shall apply Theorem A.2 to the quantities 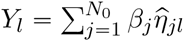. Since they are independent, we can take *D =* 1 and since, by assumption, the scaled allelic effects are bounded by 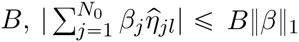 for all *l*. Then,

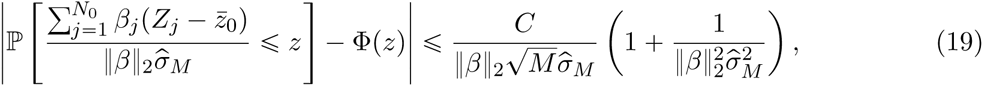

where Φ is the cumulative distribution function of a standard normal random variable and the constant *C* has an explicit expression (depending on *B* and ‖*β*‖), which can be read off from Theorem A.2. In particular, in the special case when *β*_*j*_ = 0 for all *j* ≠ *k* and *β*_*k*_ = 1, so that 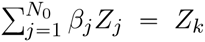, the genetic component of the trait of the *k*th individual, the constant *C* is independent of *N*_0_.

Since the vector *β* is arbitrary, this proves convergence of the joint distribution of traits in the ancestral population to a multivariate normal with the given mean vector and covariance matrix.

**C Observed traits and scaled allelic effects in the ancestral population**

When we condition on the observed trait values in our pedigree, we gain some information on the scaled allelic effect at each locus. In order to control the magnitude of this effect we use Bayes’ rule to turn it into a question about the effect of knowing the allelic effect at a given locus on the probability of observing a particular trait. We then need to be able to control

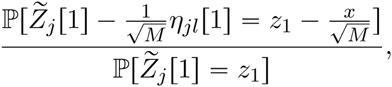

where we are interpreting the probabilities as density functions.

We begin with the case in which the parents are from the ancestral population. Using the result of Appendix B, the observed trait in each individual is, up to an error of order 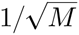, (independently) distributed according to the sum of a normal random variable with mean 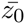 and variance 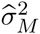 and an independent normal with mean zero and variance 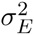.

Let us write *p(σ^2^, μ, y*) for the density function of a normally distributed random variable with mean *μ* and variance *σ*^2^. Then, taking *β*_*k*_ = 0 for *k ≠ j* and *β*_*j*_ = 1 gives

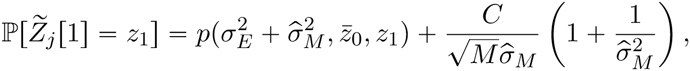

**Remark C.1** *The Central Limit Theorem of Appendix B only gives convergence of the cumulative distribution functions of the genetic component of the ancestral traits with a rate 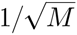. If there is a differentiable density function for each M then we can deduce the same order of convergence for the density function. If for example, allelic effects are discrete, then additional conditions would be required to approximate this ratio of probabilities by the corresponding normal distribution with this degree of accuracy as we need a* local *limit theorem to hold. McDonald (2005) surveys results in this direction. Without such conditions, the rate of convergence can be shown to be at least* 1/*M*^¼^, *but simple counterexamples show that this is optimal. Convolution with the environmental noise rescues us and gives the faster rate of convergence of densities reported here*.

The same result applied to 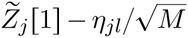, gives, up to an error of order 1/*M* which we ignore

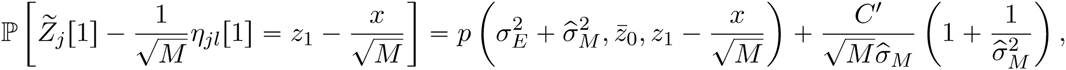

where we recall that 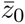 is the mean of the genetic component of the trait in generation zero. Performing a Taylor expansion of 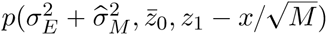 around 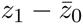 and using that

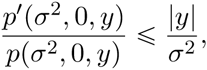

we see that for parents of individuals in the first generation,

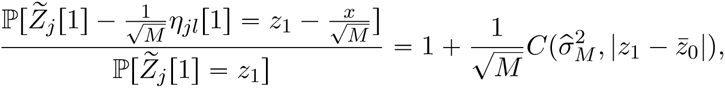

where *C*(*σ*^2^, |*z*|) was defined in equation (15). Just as with our toy model at the beginning of Section 3, we see that the approximation requires that the trait we are sampling is not ‘too extreme’.

**D One generation of reproduction**

We now establish that after one round of mating, conditional on knowing *P*^(1)^ and 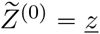,

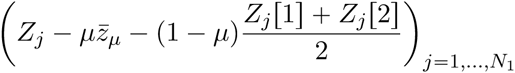

converges in distribution to a mean zero multivariate normally distributed random variable with diagonal variance-covariance matrix Σ_1_ with on-diagonal entries given by (Σ_1_)_*jj*_. The ‘remainder term’ *R*_*j*_ in (13) is given by

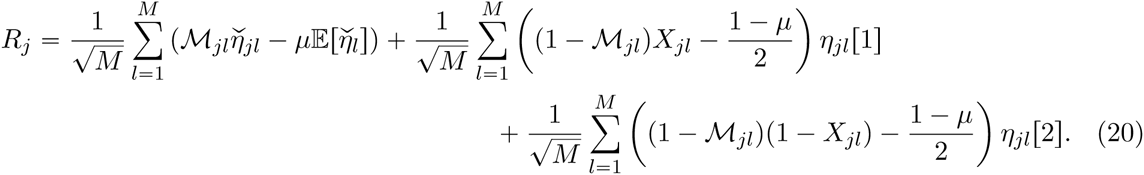

The Bernoulli random variables *M*_*jl*_ and *X*_*jl*_ are independent of both *P*^(1)^ and 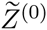 and so 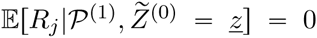. Moreover, since the Bernoulli variables in different individuals are independent, for 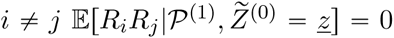. To establish 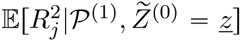, first we use Bayes’ rule to control the conditional distribution of *η*_*jl*_[1]. We condition on the whole vector of observed traits 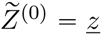, but since individuals in our ancestral population are assumed unrelated, from the perspective of *η*_*jl*_[1], this is equivalent to conditioning on the observed trait 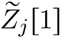 of the first parent of the *j*th individual. It is convenient to write *z*_1_ for the corresponding coordinate of ẕ.

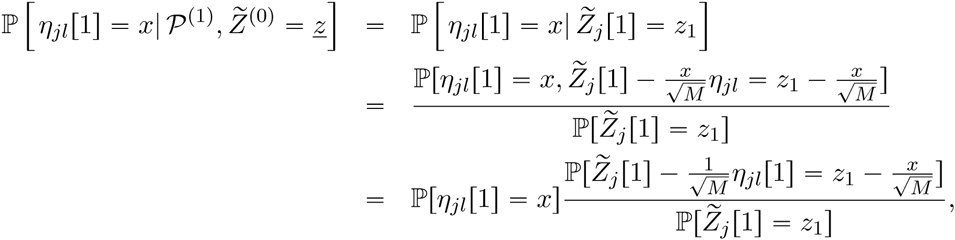

where we have used independence of inheritence at different loci and the ratio on the right should be interpreted as a ratio of probability density functions. We showed in Appendix C that the ratio in the last line is

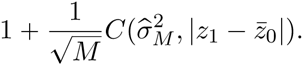

Since individuals in the ancestral population are assumed to be unrelated, *η*_*jl*_[1] and *η*_*jl*_[2] are independent and so combining the calculation above with the symmetric one for *η*_*jl*_[2] we can calculate

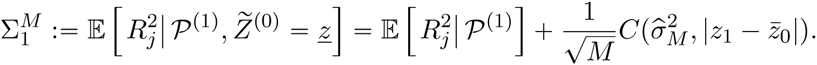

Noting that since inheritance at different loci is independent, the variance of *R*_*j*_ will be the sum of the variances at each locus, we consider the summand corresponding to a single locus, *l* say. Omitting the factor of 1/*M*, the square of the first term, corresponding to mutation, contributes

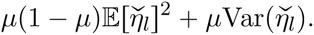

Since the variances of (1 - *M*_*jl*_.)*X*_*jl*_ and (1 - *M*_*jl*_)(1 - *X*_*jl*_) are both (1 - *μ*)(1 + *μ*)/4, the squares of the next two terms contribute

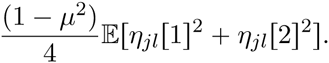

The cross terms are also non-trivial.

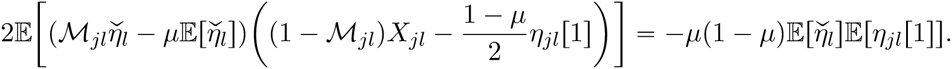

Similarly,

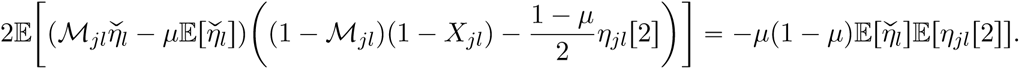

Finally,

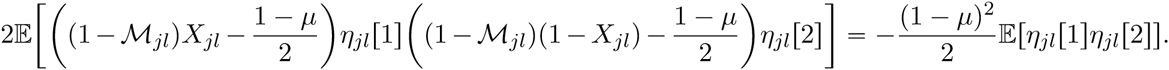

Combining these, we obtain

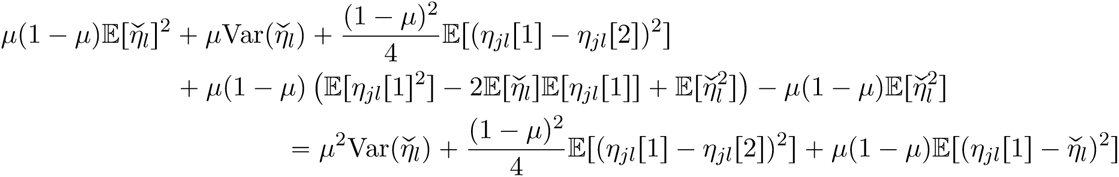

and, since *η*_*jl*_[1] is sampled from the ancestral population and so is a copy of 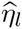 this yields

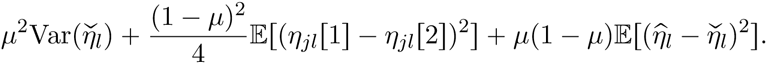

(Note that if the individual was produced by selfing, the second term is 0.) It is immediate from this calculation that the variance of our limiting distribution of traits is Σ1, as claimed. To check that the limit is a multivariate normal, we mimic what we did in the ancestral population: for an arbitrary vector *β* = (*β*_1_,…, *β*_*N*_1__) we show that 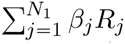 converges to a normal random variable as *M* → ∞. As before the strategy is to apply Theorem A.2. This time

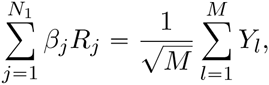

where

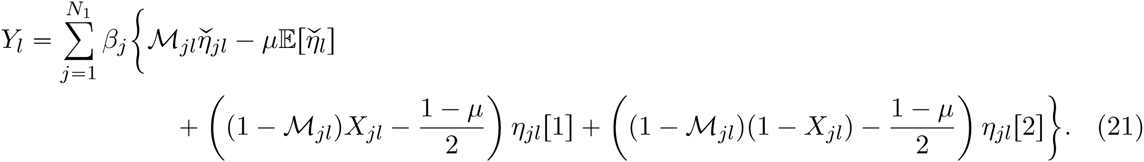

Each such term is bounded by *B*‖*β‖* and inheritance is independent at distinct loci and so Theorem A.2 yields convergence (in law) of 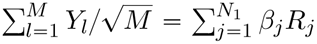 to a mean zero normal random variable with variance

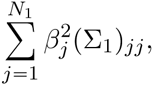

from which, since *β* was arbitrary, we deduce convergence of (*R* _1_,…, *R*_*N*_1__), conditional on knowing *P*^(1)^ (the parents of each individual in the population) and 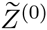 (the observed traits of all parents) to a multivariate normal with mean zero and diagonal variance-covariance matrix with on-diagonal entries identically equal to Σ1. More precisely, just as in (19),

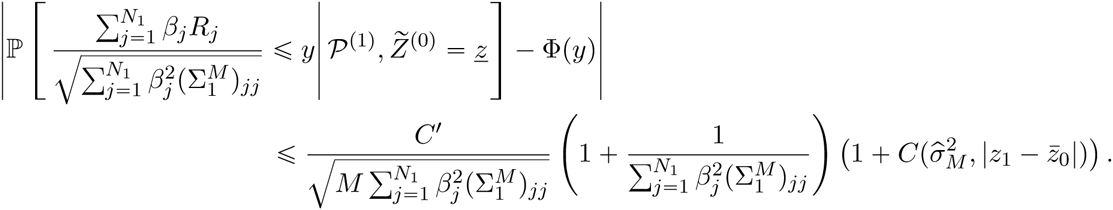

**E Generation *t***

We now provide the missing steps in the general case. We proceed by induction.

Suppose that we have proved the asymptotic normality of the vector of genetic components of trait values and (14) (that conditioning on the pedigree and the observed ancestral traits provides negligible information about the distribution of allelic types at a given locus) for all generations up to and including *(t -* 1). We have already checked generation one.

We first prove (14). Let us write *A* ~ *η*_*jl*_[1] to mean that *A* is the set of individuals in *P*^(*t-*1)^ that are identical by descent at locus *l* with the first parent of individual *j* in generation *t*. Note that *A* depends on the pedigree and the Bernoulli random variables that determine inheritance at the *l*th locus, but not on the value *η*_*jl*_[1] and so partitioning on the set *A*,

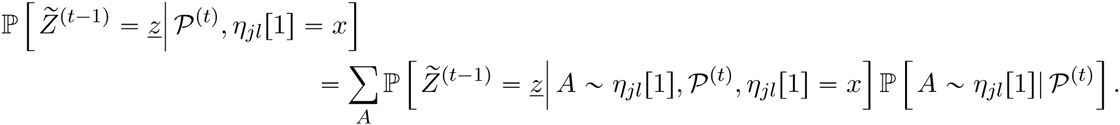

We write *a* for the eldest individual in *A* and |*a|* for the generation in which it lived. Evidently the trait values in *P*^(*t*-1)^\*A* do not depend on *η*_*jl*_[1]. Moreover, if we further partition on the value of *Z*_*a*_ (the genetic component of the trait of the eldest member of *A*), we see that for all *a′* ∈ *A\a*, the probability that 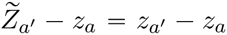 is independent of the value of *η*_*jl*_[1]. In other words, the dependence of the trait values in the pedigree on *η*_*jl*_[1] is entirely captured by

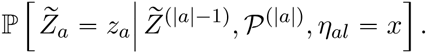

Since *a* lives at the latest in generation *t -* 1, we can use our inductive hypothesis to write that

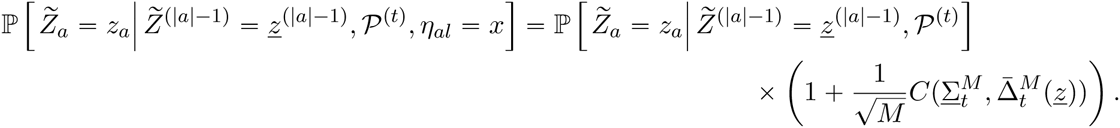

Since we have successfully eliminated all the conditioning on the value of *η*_*jl*_[1], we can now rearrange our calculations to give Equation (14) and

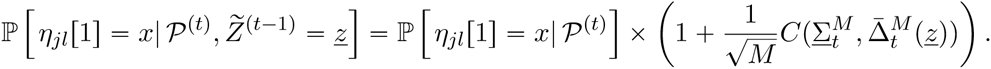

We can perform entirely analogous calculations for the joint law of *η*_*jl*_[1] and *η*_*jl*_[2].

Now consider the mean zero random variable *R*_*j*_. That 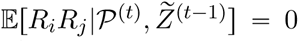 for *i* ≠ *j* follows exactly as before and the calculation of 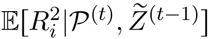 also proceeds almost exactly as for generation one. The only distinction is that

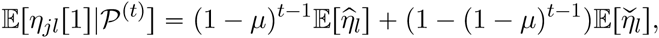

and similarly

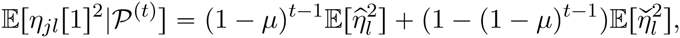

and so the contribution to 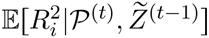 from the *l*th locus becomes

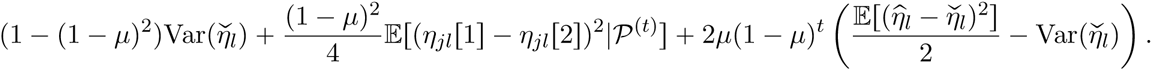

Summing over loci yields (11).

Now, exactly as we did for generation one, we can fix *β*_1_,…, *β*_*N*_*t*__ ∈ ℝ and apply Theorem A.2 to *Y*_*t*_ given by (21) (with the same bound) to deduce that conditional on *P*^(*t*)^ and 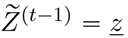,

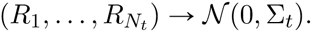

In particular,

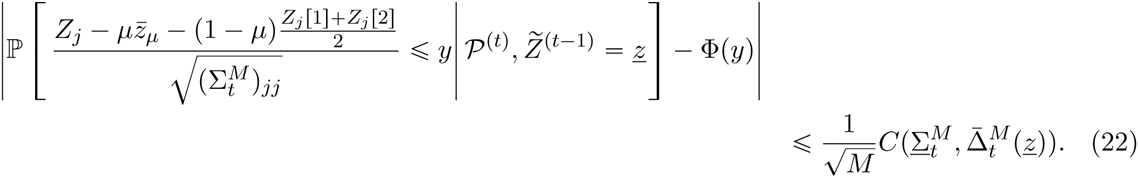

**F Environmental noise: conditioning multivariate Gaussian vectors**

In order to estimate the proportion of an observed trait that is due to environmental noise, and thus make predictions about offspring traits, we need a standard result for conditioning multivariate normal random vectors on their marginal values which, for ease of reference, we record here.

**Theorem F.1** *Suppose that*

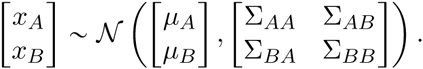

*Then*

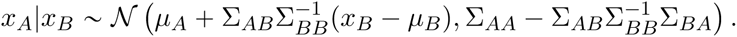

The proof can be found e.g. in Brockwell & Davis (1996) (Prop. 1.3.1 in Appendix A). To see how this leads to a recurrence for the conditional values of the environmental noise, we begin with generation zero. In this case there are just two components to consider, (*R*_*j*_)_*j*_,…, _*N*_0__ and (*E*_*j*_)_*j* = 1,…, *N*_0__, each of which is (at least asymptotically) a mean zero Gaussian with diagonal variance-covariance matrix. We wish to calculate *x*_*A*_|*x*_*B*_ where *x*_*A*_ = (*E*_*j*_)_*j* = 1,…, *N*_0__ and 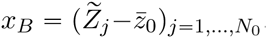. We have

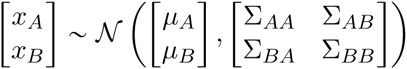

where 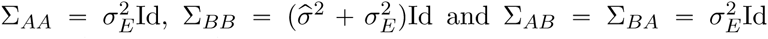. Applying Theorem F.1, 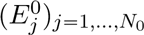 (as *M* → ∞)is a Gaussian random variable with mean vector

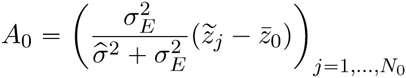

and variance-covariance matrix

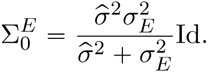

For the recursive step, we now set

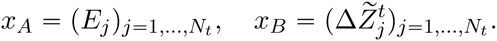

Then, conditionally on *P*^(*t*)^ and 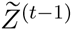,

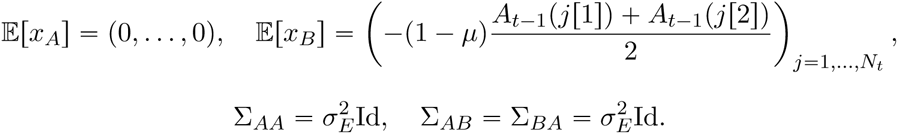

The more complex term is

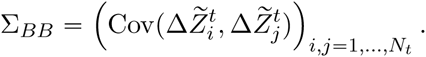

We treat the cases *i = j* and *i* ≠ *j* separately. In the expression below, *i*[*a*] is the *a*th parent of individual *i* (with *a* ∈ {1,2}). First suppose that *i* ≠ *j*, again conditionally on *P*^(*t*)^ and 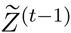,

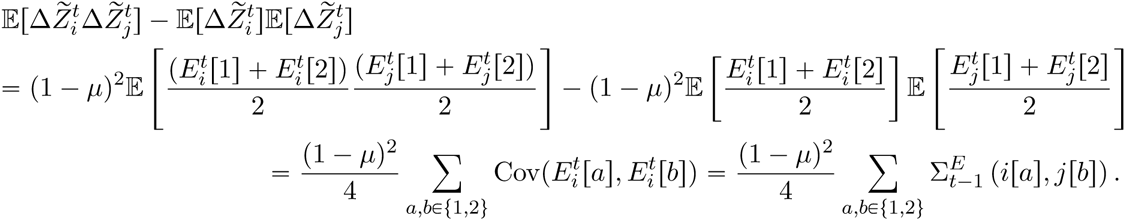

If *i* = *j*,

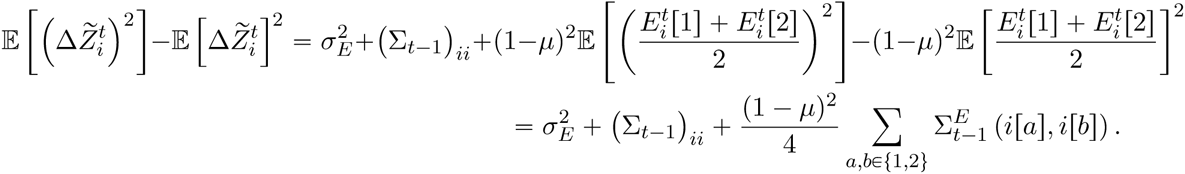

Again applying Theorem F.1, we obtain that *E*^*t*^ has mean vector

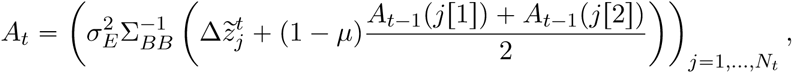

and variance-covariance matrix

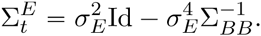

**G A coupling argument**

We do not spell out all of the details of the proof of convergence to a multivariate normal in the presence of epistasis. However, we illustrate a useful coupling argument by explaining how to use it to prove that in generation one, *𝔼[*∆*Z*_*j*_] is of order 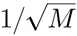. Recall that

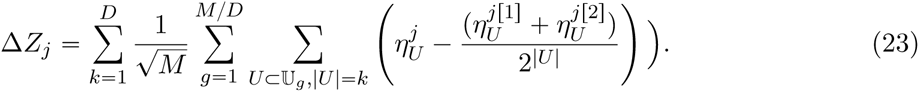

We use the notation 1_*U←j*[1]_ for the Bernoulli random variable which takes the value 1 when all the loci in *U* were inherited from *j*[1]. Then

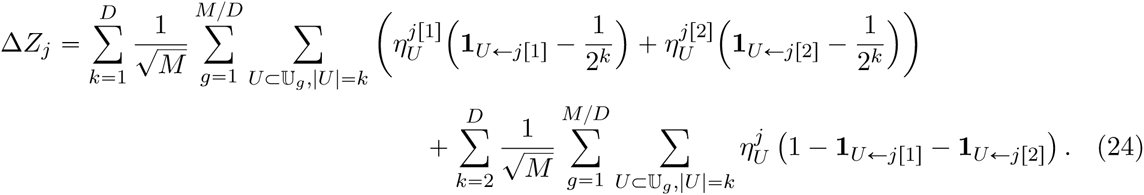

Notice that the second sum starts from *k =* 2. Because the Bernoulli random variables that determine inheritance are independent of the parental allelic effects, the expectation of the first sum is zero. A priori, the second term could be order one, but we now argue that it is order 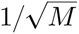. The idea is a simple coupling argument. The analogue of (14) tells us that conditioned on the values *z*_*A*_, … *z*_*A(k*)_ in the ancestral population, the allelic state of the parents at the loci in *𝕌*_*g*_ have the original distribution 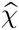 with probability 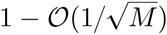. Because the state space of *(χ*_*l*_)_𝕌_*g*__ is finite, this means that we can couple the conditioned distributions of the allelic states for loci *l* ∈ *𝕌*_*g*_ of each parent in such a way that with probability 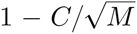 they are drawn from the unbiased distribution 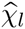 and with probability 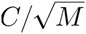 they are drawn from some modified distribution 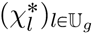 which is independent of 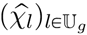. As a result, summing over all admissible inheritance patterns, we obtain

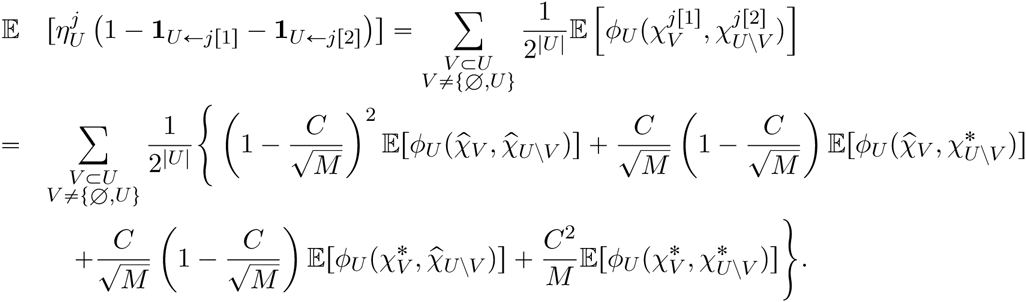

The first term is zero by assumption and condition (17) guarantees that so are the second and third terms. In total this sum is at most *C*_*U*_/*M* where *C*_*U*_ is bounded uniformly in *U* (since the allelic effects are) and so multiplying by 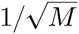 and summing over *k* and *g* we find that 𝔼[∆*Z*_*j*_] is order 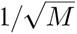 as claimed.

## Notes

* NHB supported in part by ERC Grant 250152

† AME supported in part by EPSRC Grant EP/K034316/1

‡ AV supported by the chaire Modélisation Mathématique et Biodiversité of Veolia Environment - École Polytechnique - Museum National d’Histoire Naturelle - Fondation X

## References

Baldi, P. and Rinott, Y. (1989). On normal approximations of distributions in terms of dependency graphs. Ann. Probab., 17:1646–1650.

Barton, N. H. (2010). What role does natural selection play in speciation? Phil. Trans. Roy. Soc. London B, 365:1825–1840.

Barton, N. H. and Etheridge, A. M. (2011). The relation between reproductive value and genetic contribution. Genetics, 188:953–973.

Barton, N. H. and Keightley, P. D. (2002). Understanding quantitative genetic variation. Nature Reviews Genetics, 3(1):11–21.

Barton, N. H. and Shpak, M. (2000). The stability of symmetrical solutions to polygenic models. Theor. Popul. Biol., 57:249–264.

Barton, N. H. and Turelli, M. (2004). Effects of genetic drift on variance components under a general model of epistasis. Evolution, 58:2111–2132.

Brockwell, P. J. and Davis, R. A. (1996). Introduction to time series and forecasting. Springer texts in statistics.

Bulmer, M. G. (1971). The effect of selection on genetic variability. American Naturalist, 105:201–211.

Bulmer, M. G. (1974). Linkage disequilibrium and genetic variability. Genet. Res., 23:281–289.

Bulmer, M. G. (1980). The mathematical theory of quantitative genetics. Oxford University Press.

Bulmer, M. G. (1998). Galton’s law of ancestral heredity. Heredity, 81:579–585.

Bulmer, M. G. (2004). Did Jenkin’s swamping argument invalidate Darwin’s theory of natural selection? British Journal of the History of Science, 37:1–17.

Cavalli-Sforza, L. L. and Bodmer, W. F. (1971). The genetics of Human populations. Freeman, San Francisco.

Charlesworth, B. (2015). Causes of natural variation in fitness: Evidence from studies of Drosophila populations. Proc. Nat. Acad. Sci. U.S.A., 112(6):1662–1669.

Cockerham, C. C. (1954). An extension of the concept of partitioning hereditary variance for analysis of covariances among relatives when epistasis is present. Genetics, 39:859–882.

Cooley, J. W. and Tukey, J. W. (1965). An algorithm for the machine calculation of complex Fourier series. Math. Comput., 19:297–301.

Davis, A. S. (1871). The North British Review and the Origin of Species. Nature, 5:161.

Diehl, S. R. and Bush, G. L. (1989). The role of habitat preference in adaptation and speciation. In Speciation and its consequences, pages 345–365. Sinauer Press, Sunderland Mass.

Doebeli, M. (1996). A quantitative genetic competition model for sympatric speciation. J. Evol. Biol., 9:893–910.

Fisher, R. A. (1918). The correlation between relatives on the supposition of Mendelian inheritance. Proc. Roy. Soc. Edinburgh, 52:399–433.

Frankham, R. (1996). Relationship of genetic variation to population size in wildlife. Conserv. Biol., 10:1500–1508.

Galton, F. (1877). Typical laws of heredity. Nature, 15:492–495, 512-514, 532-533.

Galton, F. (1885). Regression towards normality in hereditary stature. Journal of the Anthropological Institute, 15:246–263.

Galton, F. (1889). Natural Inheritance. Macmillan, London.

Gauss, C. F. (1886). Theoria interpolationis methodo nova tractata Werke band 3, 265–327. Göttingen: Königliche Gesellschaft der Wissenschaften.

Good, B. H. and Desai, M. M. (2014). Deleterious passengers in adapting populations. Genetics, 198:1183–1208.

Haldane, J. B. S. (1931). A mathematical theory of natural and artificial selection. VII. Selection intensity as a function of mortality rate. Proc. Camb. Phil. Soc., 27:131–136.

Henderson, C. R. (2014). Estimation of variance and covariance components. Biometrics, 9:226–252.

Hill, W. G. (1982). Rates of change in quantitative traits from fixation of new mutations. Proc. Nat. Acad. Sci. U.S.A., 79:142–145.

Hill, W. G. (2014). Applications of population genetics to animal breeding, from wright, fisher and lush to genomic prediction. Genetics, 196:1–16.

Hill, W. G. and Kirkpatrick, M. (2010). What animal breeding has taught us about evolution. Annu. Rev. Ecol. Evol. Syst., 41:1–21.

Jenkin, F. (1867). Origin of Species. North British Review, 46:277–318.

Johnson, T. and Barton, N. H. (2005). Theoretical models of selection and mutation on quantitative traits. Phil. Trans. Roy. Soc. London B, 360:1411–1425.

Kempthorne, O. (1954). The correlation between relatives in a random mating population. Proc. Roy. Soc. London B, 143:102–113.

Kimura, M. (1965). A stochastic model concerning the maintenance of genetic variability in quantitative characters. Proc. Nat. Acad. Sci. U.S.A., 54:731–736.

Kondrashov, A. S. (1984). On the intensity of selection for reproductive isolation at the beginnings of sympatric speciation. Genetika, 20:408–415.

Kruuk, L. E. B. (2004). Estimating genetic parameters in natural populations using the “animal model”. Phil. Trans. Roy. Soc. London B, 359:873–890.

Lande, R. (1976a). The maintenance of genetic variability by mutation in a polygenic character with linked loci. Genet. Res., 26:221–235.

Lande, R. (1976b). Natural selection and random genetic drift in phenotypic evolution. Evolution, 30:314–334.

Lande, R. and Arnold, S. J. (1983). The measurement of selection on correlated characters. Evolution, 37:1210–1226.

Lange, K. (1978). Central limit theorems for pedigrees. J. Math. Biol., 6:59–66.

Loewe, L. and Charlesworth, B. (2006). Inferring the distribution of mutational effects on fitness in drosophila. Biology Letters, 2(2):426–430.

Lush, J. L. (1937). Animal breeding plans. Ames, Iowa.

Lynch, M. and Walsh, B. (1997). Genetics and analysis of quantitative traits. Sinauer Press.

Magnello, M. E. (1998). Karl Pearson’s mathematization of inheritance: from ancestral heredity to Mendelian genetics. Annals of Science, 55:35–94.

McDonald, D. R. (2005). The local limit theorem: a historical perspective. JIRSS, 4(2):73–86.

Meuwissen, T., Hayes, B., and Goddard, M. (2013). Accelerating improvement of livestock with genomic selection. Annu. Rev. Anim. Biosci., 1(1):221–237.

Pearson, K. (1896). Mathematical contributions to the theory of evolution. III. Regression, heredity, and panmixia. Phil. Trans. Roy. Soc. London A, pages 1–67.

Pearson, K. (1897). Mathematical contributions to the theory of evolution. On the law of ancestral heredity. Proc. Roy. Soc. London, 62:386–412.

Pearson, K. (1904a). Mathematical contributions to the theory of evolution. XII. On the generalised theory of alternative inheritance with special reference to Mendel’s Laws. Phil. Trans. Roy. Soc. London A, 203:53–86.

Pearson, K. (1904b). A Mendelian view of the law of ancestral inheritance. Biometrika, 3:109–112.

Pearson, K. (1909). The theory of ancestral contributions of a Mendelian population mating at random. Proc. Roy. Soc. London, 81:225–229.

Polechova, J. and Barton, N. H. (2005). Speciation through competition: a critical review. Evolution, 59:1194–1210.

Provine, W. (1971). The origins of theoretical population genetics. Univ. of Chicago Press. Chicago.

Rinott, Y. (1994). On normal approximation rates for certain sums of dependent random variables. J. Comp. Appl. Math., 55:135–143.

Robertson, A. (1960). A theory of limits in artificial selection. Proc. Roy. Soc. London B, 153:234–249.

Robertson, A. (1961). Inbreeding in artificial selection programmes. Genet. Res., 2:189–194.

Robertson, A. (1966). A mathematical model of the culling process in dairy cattle. Animal Production, 8:95–108.

Santiago, E. (1998). Linkage and the maintenance of variation for quantitative traits by mutation-selection balance: an infinitesimal model. Genet. Res., pages 161–170.

Turelli, M. (1984). Heritable genetic variation via mutation-selection balance: Lerch’s Zeta meets the abdominal bristle. Theor. Popul. Biol., 25:1–56.

Turelli, M. and Barton, N. H. (1994). Statistical analyses of strong selection on polygenic traits: What, me normal? Genetics, pages 1–29.

Weber, K. E. and Diggins, L. T. (1990). Increased selection response in larger populations. II. Selection for ethanol vapour resistance in Drosophila melanogaster at two population sizes. Genetics, 125:585–597.

Weissman, D. B. and Barton, N. H. (2012). Limits to the rate of adaptation in sexual populations. PloS Genetics, 8(6):e1002740. doi:10.1371/journal.pgen.1002740.

Willi, Y., van Buskirk, J., and Hoffmann, A. A. (2006). Limits to the adaptive potential of small populations. Ann. Rev. Ecol. Evol. Syst., 37:433–458.

Wray, N. R. (1990). Accounting for mutation effects in the additive genetic variance-covariance matrix and its inverse. Biometrics, pages 177–186.

Wright, S. (1935). The analysis of variance and the correlation between relatives with respect to deviations from an optimum. J. Genet., 30:243–256.

Yang, J., Benyamin, B., McEvoy, B. P., S, S. G., Henders, A. K., Nyholt, D. R., Madden, P. A., Heath, A. C., Martin, N. G., Montgomery, G. W., Goddard, M. E., and Visscher, P. M. (2010). Common SNPs explain a large proportion of the heritability for human height. Nature Genet., 42:565–569.

Yule, G. U. (1902). Mendel’s Laws and their probable relations to inter-racial heredity. New Phytologist, 1:193-207, 221-237.

Yule, G. U. (1907). On the theory of inheritance of quantitative compound characters on the basis of Mendel’s laws. In Report of the Conference on Genetics. Royal Horticultural Soc.

